# Effect of mitochondrial circulation on mitochondrial age density distribution

**DOI:** 10.1101/2022.12.01.518783

**Authors:** Ivan A. Kuznetsov, Andrey V. Kuznetsov

## Abstract

Recent publications report that although the mitochondria population in an axon can be quickly replaced by a combination of retrograde and anterograde axonal transport (often within less than 24 hours), the axon contains much older mitochondria. This suggests that not all mitochondria that reach the soma are degraded and that some are recirculating back into the axon. To explain this, we developed a model that simulates mitochondria distribution when a portion of mitochondria that return to the soma are redirected back to the axon rather than being destroyed in somatic lysosomes. Utilizing the developed model, we studied how the percentage of returning mitochondria affects the mean age and age density distributions of mitochondria at different distances from the soma. We also investigated whether turning off the mitochondrial anchoring switch can reduce the mean age of mitochondria. For this purpose, we studied the effect of reducing the value of a parameter that characterizes the probability of mitochondria transition to the stationary (anchored) state. The reduction in mitochondria mean age observed when the anchoring probability is reduced suggests that some injured neurons may be saved if the percentage of stationary mitochondria is decreased. The replacement of possibly damaged stationary mitochondria with newly synthesized ones may restore the energy supply in an injured axon. We also performed a sensitivity study of the mean age of stationary mitochondria to the parameter that determines what portion of mitochondria re-enter the axon and the parameter that determines the probability of mitochondria transition to the stationary state.

## 1. Introduction

The primary purpose of mitochondria in the cells is to generate easily accessible chemical energy in the form of ATP. In addition, mitochondria serve various other functions, such as calcium buffering. They are also known for their involvement in apoptosis [1-3], particularly in the context of stroke-induced injury [4].

Since mitochondrial proteins have a limited half-life, these proteins (or the entire mitochondrion) need to be periodically replaced [5]. Old and damaged mitochondria are usually degraded in somatic lysosomes [6,7]. Mitochondria are continuously transported in axons [8]. The representative length of mitochondria is between 0.5 and 10 μm [9]. Hence, mitochondria are too large to exhibit any significant diffusivity. Mitochondria transport is accomplished by molecular motors, kinesin-1 (and possibly kinesin-3) in the anterograde direction and by cytoplasmic dynein in the retrograde direction [10,11]. Mitochondria move with an average velocity of 0.4-0.8 μm/s [12]. To supply energy to energy demand sites in the axon, mitochondria can dock near these demand sites [13-15].

In neurons, most of the degradation of old and damaged mitochondria occurs in the soma. It was previously hypothesized that mitochondria are transported to the soma for final degradation [16-18]. However, ref. [19] reported that after retrogradely transported mitochondria depart from the axon terminal, a portion of them remain in the axon for days. The persistence of mitochondria after leaving the terminal for a prolonged duration may indicate that they will never reach the soma, as they spend much time in the stationary state. Alternatively, this may suggest that retrogradely transported mitochondria undergo recycling in the soma. Based on this evidence, we hypothesize that not all mitochondria that enter the soma from the axon undergoing retrograde movement are degraded in the soma; a portion return to the axon. This hypothesis is supported by unpublished observations reported in ref. [3]. These observations suggest that retrogradely moving mitochondria, which are traditionally expected to be segregated for degradation through mitophagy upon entering the soma, instead merge with the existing mitochondria in the soma. This fusion leads to the exchanging of components between the returning mitochondria and the newly synthesized mitochondria in the soma, producing repaired mitochondria that are ready to return to the axon.

A high level of energy is required for neurons to survive an injury. Since the distance to which ATP can be transported by diffusion is limited, replacing damaged mitochondria with new ones is critical for neuron survival [20]. Axonal regeneration after injury or ischemic stress [21] can be promoted by turning off mitochondria anchoring. This leads to faster replacement of damaged mitochondria with healthy mitochondria [22]. Our model simulates the reduction of mitochondrial anchoring by decreasing a value of the parameter that characterizes the probability of mitochondrial anchoring, *p*_*s*_.

The return of mitochondria to the axon must be associated with changing the type of molecular motors that move the mitochondria. Since mitochondria are driven toward the soma by dynein motors, to turn around and re-enter the axon they must switch from dynein motors to kinesin motors.

Ref. [23] used mean-field equations to study the delivery of motor-cargo complexes to branched axons. A quantity called mitochondrial health was introduced in ref. [24] and treated as a proxy for mitochondrial membrane potential. Models that treat mitochondrial health as a decaying component were developed in ref. [25].

This paper adopts a different approach by using a compartment-based model. Compartments do not necessarily need to be physical volumes with actual bottlenecks between them. A compartmental model, for example, could be used to simulate a tracer in tissue, which binds to various specific target receptor sites [26,27]. In this paper we need to simulate mitochondria transport between various mitochondrial demand sites. Each demand site is represented by three compartments containing populations of anterograde, stationary, and retrograde mitochondria. We developed this model in our previous publications and utilized it for determining the mean age of mitochondria along the axon length [28] and for computing the mean age and age density distributions of mitochondria in a branching axon [29].

The main question that we ask in this paper is how the mean age and age density distributions of mitochondria in the compartments located at different distances from the soma are affected by some of the mitochondria that exit the axon changing the motors that propel them to kinesin and returning to the circulation. This would explain the presence of older mitochondria in the axon, reported, for example, in ref. [19]. Intuitively, this should increase the mean age of mitochondria in the axon as a result of the described process in which older mitochondria re-enter the axon. Another question we attempt to answer in this paper is whether making mitochondria less likely to stop would contribute to a faster renewal of mitochondria (which in our model will appear as a reduction of the mean age of mitochondria). This would improve chances for neuronal recovery after ischemic stress or injury [22].

## 2. Materials and models

### 2.1 Problem setup

Mitochondria usually dock near energy demand sites in the axon [30]. We simulated the axon as consisting of *N* demand sites (Fig. 1). Mitochondria in the axon can be divided into stationary, anterograde, and retrograde, with transitions possible between these three pools [13]. We therefore represented each demand site as that consisting of three compartments which contain stationary, anterograde, and retrograde mitochondria. The total number of compartments in our multi-compartment model [31-33] is thus 3*N*, see Fig. 2 (we used *N*=10). Following ref. [25], we define the demand sites as narrow sections of the axon where stationary mitochondria are localized. The selection of the number of the demand sites should be interpreted as a means of simulating axonal morphology rather than discretization. It is important to acknowledge that in a real axon, the number of the demand sites will be substantially larger than 10. A narrow demand site is assumed to be located at the center of each compartment, and the compartment is assumed to surround it.

**Fig. 1.**
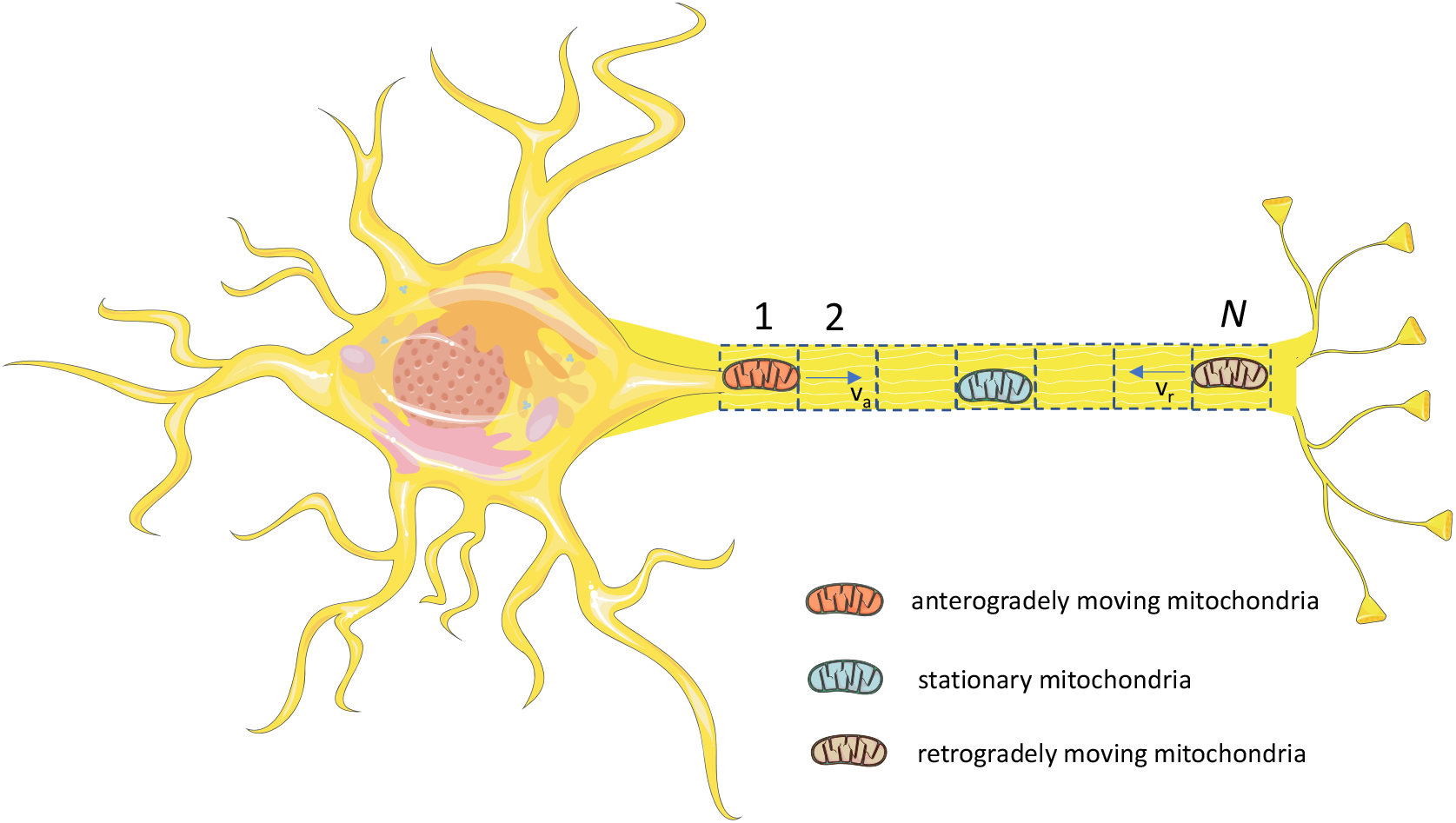
(a) Schematic diagram of a neuron with an axon that is assumed to contain *N* energy demand sites. Demand sites are numbered starting with the most proximal (site 1) to the most distal (site *N*). Figure is generated with the aid of Servier Medical Art, licensed under a creative commons attribution 3.0 generic license, http://Smart.servier.com.

**Fig. 2.**
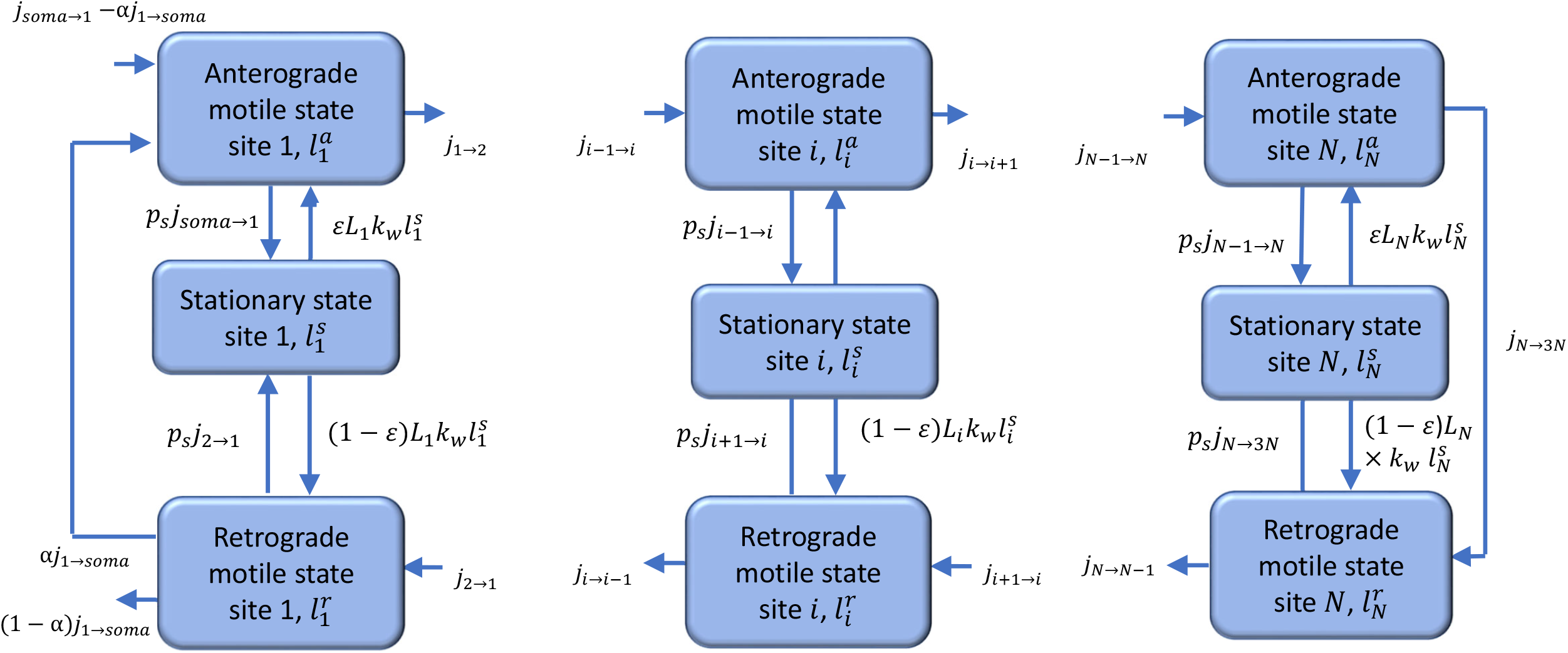
A diagram of a compartmental model showing transport in the compartments surrounding the demand sites in the axon. Arrows show mitochondria exchange between the transiting (anterograde and retrograde) states in adjacent demand sites, mitochondria capture into the stationary state and re-release from the stationary state. A portion of mitochondria re-enter the axon, and the rest return to the soma for degradation. Newly synthesized mitochondria entering the axon from the soma are assumed to have zero age.

The primary motivation for employing a multi-compartment model is our desire to use the methodology for computing the particle age density distribution in the compartments developed in refs. [34,35]. We used a similar approach to simulate transport of dense core vesicles in *Drosophila* motoneurons [36,37]. The demand sites in the axon terminal were numbered from the most proximal (site 1) to the most distal (site *N*) (Fig. 1).

To characterize mitochondria concentration in the axon, we used the total length of mitochondria in a particular kinetic state (Fig. 2) per unit axon length, measured in (μm of mitochondria)/μm. The dependent variables utilized in the model are given in Table S1.

### 2.2 Governing equations and the model for the mean age and age density distributions of mitochondria in the demand sites

The total length of mitochondria is a conserved quantity. The equations stating the conservation of the total length of mitochondria in 3*N* compartments (Fig. 2) are stated using the methodology developed in ref. [31]. In a vector form, these equations are

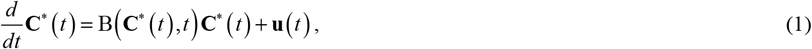

where *t* is the time and **C**^∗^is the state vector, following the terminology used in ref. [34]. **C**^∗^is a column vector; its components are defined as follows:

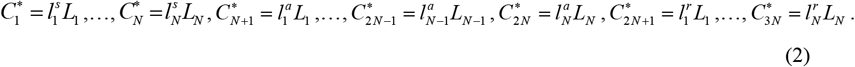

In Eq. (2)

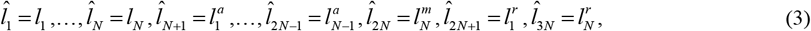

where *l*_*i*_ is the total length of mitochondria per unit length of the axon in stationary, anterograde, or retrograde states in the compartment by the *i*th demand site (Table S1).

Model parameters are summarized in Table S2. In Eq. (2) *L*_*i*_ (*i*=1,…,*N*) is the length of an axonal compartment around the *i*th demand site. Since the axonal length *L*_*ax*_ was split into *N* compartments of equal length, *L*_*i*_ = *L*_*ax*_ / *N*.

The first *N* components of the state vector represent the total length of mitochondria in stationary states, the next *N* components (*N*+1,…,2*N*) represent the total length of mitochondria in anterograde states, and the last *N* components (2*N*+1,…,3*N*) represent the total length of mitochondria in retrograde states. The definition of vector **u**(*t*) is given below in Eqs. (45) and (46).

Matrix B(3*N*,3*N*) is composed following ref. [31]. It accounts for the internal mitochondria fluxes between the compartments, the external mitochondria flux entering the axon from the soma, and the mitochondria flux leaving the axon, part of which returns to the soma for degradation and part re-enters the axon (Fig. 2). The analysis of mitochondria fluxes to and from the stationary, anterograde, and retrograde compartments surrounding the most proximal demand site gives equations for the following elements of matrix B:

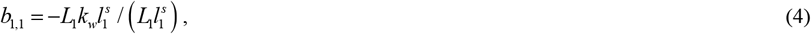

where *k*_*w*_ is the kinetic constant characterizing the rate of mitochondria re-release from the stationary state into the mobile states.

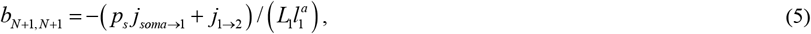

where *j* denotes the flux of mitochondria length between the compartments and *p*_*s*_ is the probability of mitochondria transitioning from a moving to the stationary state.

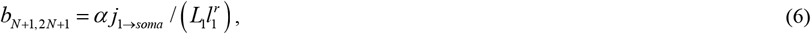

where *j*_*soma*⟶1_ is the total flux of anterograde mitochondria entering the most proximal demand site, which consists of mitochondria newly synthesized in the soma and mitochondria returning from their journey in the axon. Our model assumes that a portion of mitochondrial flux exiting the axon, *α j*_1⟶*soma*_, returns to the axon while the other portion of the flux, (1 − *α*) *j*_1⟶*soma*_, is degraded in somatic lysosomes.

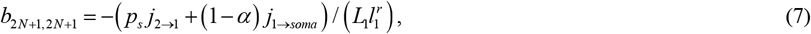

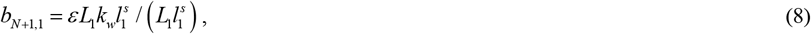

where *ε* is the portion of mitochondria that re-enter the anterograde component and (1 − *ε*) is the portion of mitochondria that re-enter the retrograde component (Fig. 2, Table S2).

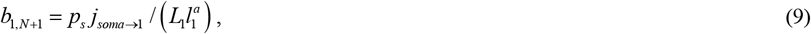

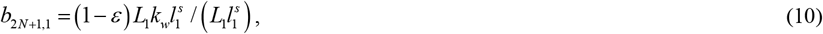

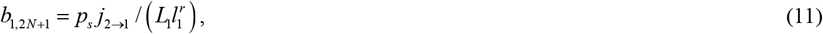

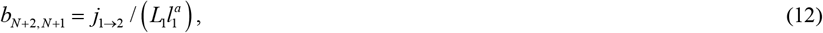

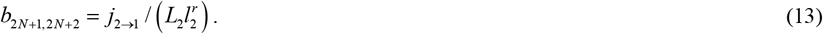

By analyzing mitochondria fluxes to and from the compartments surrounding demand site *i* (*i*=2,…,*N*-1), equations for the following elements of matrix B are obtained:

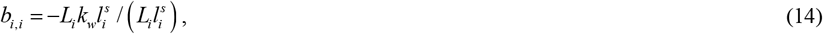

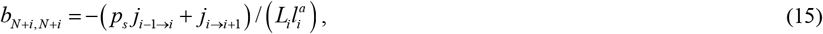

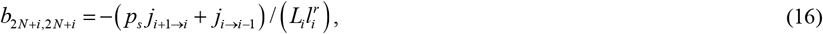

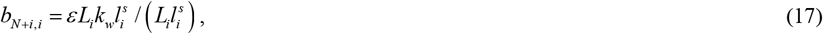

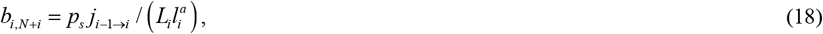

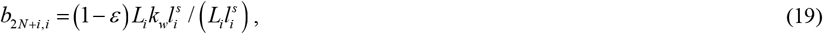

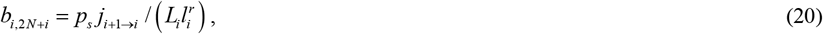

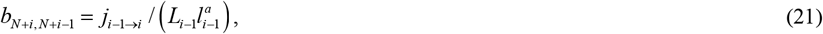

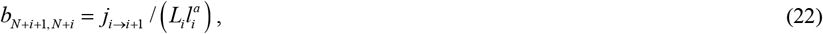

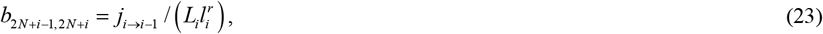

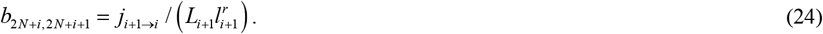

The analysis of mitochondria fluxes to and from the compartments surrounding the most distal demand site gives equations for the following elements of matrix B:

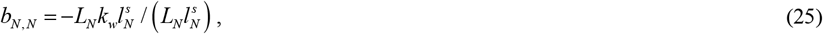

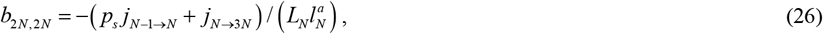

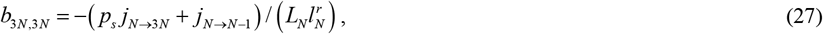

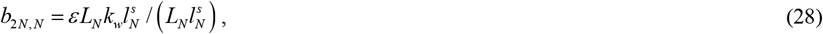

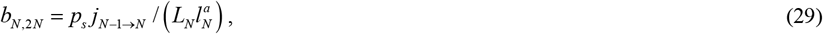

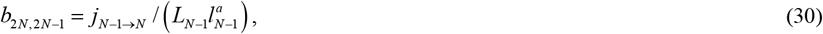

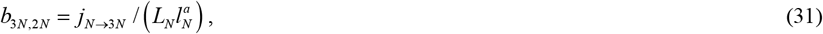

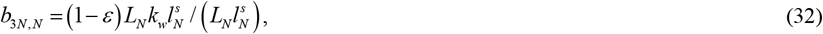

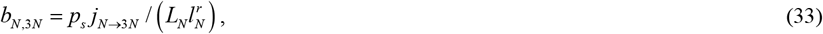

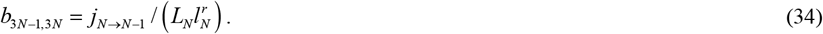

Other elements of matrix B, except for those given by Eqs. (4)-(34), are equal to zero.

Eqs. (4)-(34) were supplemented by the following equations that simulate fluxes of mobile mitochondria between the demand sites. Equations for anterograde fluxes (Fig. 2) are given by the following equations:

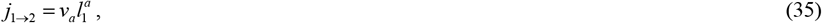

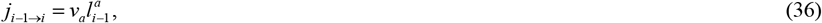

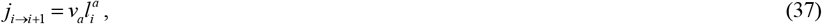

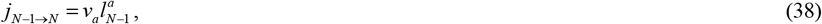

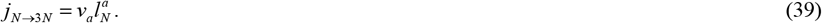

Equations for retrograde fluxes (Fig. 2) are given by the following equations:

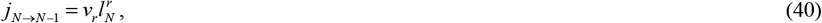

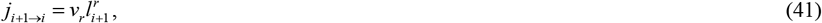

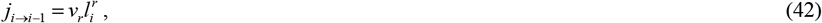

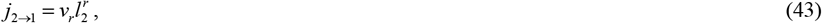

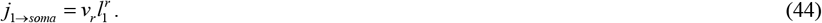

The *N*+1^th^ element of column vector **u** is given by the following equation:

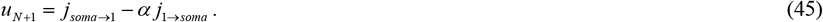

In Eq. (45) *j*_*soma*⟶1_ is the total flux of anterograde mitochondria entering the most proximal demand site. It consists of mitochondria newly synthesized in the soma and mitochondria that just left the axon and re-enter the axon anterogradely. Eq. (45) means that the flux of newly synthesized mitochondria entering the axon from the soma, *u*_*N* +1_, is adjusted in our model depending on the flux of mitochondria that re-entered the axon after leaving it retrogradely, *α j*_1⟶*soma*_. This is done to keep the total flux of mitochondria entering the axon (newly synthesized plus returning) constant and independent of *α*.

The only flux entering the compartmental system displayed in Fig. 2 is the flux of newly synthesized mitochondria entering the *N*+1^th^ compartment. Therefore, all other elements of vector **u** are equal to zero:

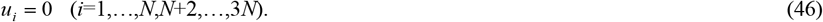

The aim of using Eqs. (1)-(46) was to utilize advanced methods developed for compartmental systems [34,35,38,39] to analyze the average age and distribution of age density of mitochondria in demand sites at varying distances from the soma. A different approach to expressing the conservation equations for the overall length of mitochondria in the axon, as presented in ref. [28], is described in section S1 of Supplemental Materials. Equations in section S1 directly state mitochondria conservation in the demand site, rather than combining stationary, anterograde, and retrograde states into a single vector. Consequently, they are easier to interpret physically. However, they are not suitable for calculating the mean age and age density distributions of mitochondria, as those require stating the conservation equations in the vector form given by Eq. (1).

We used the formulas reported in ref. [35] to obtain results for the steady-state situation (when the left-hand side of Eq. (1) is equal to zero). The solution of Eq. (1) for a steady-state situation is given by the following equation:

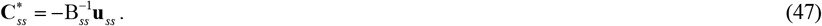

In Eq. (47), the superscript –1 on a matrix denotes the inverse of the matrix and the subscript *ss* denotes steady-state.

The mean ages and age density distributions of mitochondria in various demand sites were computed using the method described in refs. [34,35]. The mean ages of mitochondria in various compartments displayed in Fig. 2 at steady-state are found from the following equation:

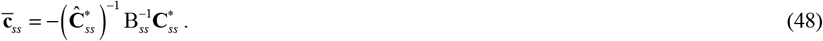

In Eq. (48),

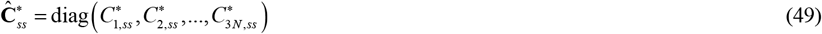

is the diagonal matrix with the components of the state vector, defined in Eq. (2), on the main diagonal. These components (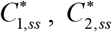, etc.) are calculated at steady-state. The left-hand side of Eq. (48) is a column vector composed of the mean ages of mitochondria in various compartments:

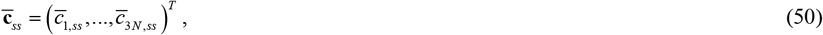

where superscript *T* denotes the transposed matrix.

The age densities of mitochondria in various compartments are given by the elements of the following vector:

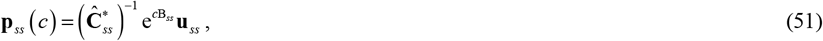

where e is the matrix exponential.

Eqs. (47)-(51) allow finding steady-state solutions directly. We implemented these equations using standard MATLAB operators, such as matrix inverse and matrix exponential.

We defined the following vectors, each of size *N*, which represent the mean ages of mitochondria in the stationary, anterograde, and retrograde states in the demand sites, respectively:

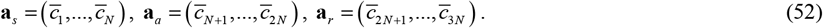

## 3. Results

### 3.1 Total length of mitochondria per unit axon length in the demand sites and mitochondria fluxes between the demand sites

Model parameters were chosen using the computational study of ref. [25], which found parameter values that optimized mitochondrial health. The parameter values used in our research are summarized in Table S2.

For *α* = 0, *p*_*s*_ = 0.4, the distribution of the total length of mitochondria per unit length of the axon is uniform along the length of the axon (Fig. S1a). The uniform distribution is expected for all parameters, regardless of *α*. This is because the model assumes that no mitochondria are destroyed while in the axon, although (1 − *α*) portion of all mitochondria are assumed to be destroyed in the soma when they re-enter the soma from the axon. The uniform distribution displayed in Fig. S1a is consistent with ref. [40] which noted that in axons the mitochondrial density is independent of the distance from the soma (Fig. 1C in ref. [40]). A constant mitochondrial density of 5 mitochondria/100 μm was also reported for growing neuronal processes in *C. elegans* [41].

The return of 30% (*α* = 0.3, *p*_*s*_ = 0.4) of exiting mitochondria back to the axon results in the same uniform distribution of mitochondria along the axon length (Fig. S1a). This is because we assumed that the total flux of mitochondria entering the first demand site is the same for all cases computed in this paper. The flux is independent of what portion of returning mitochondria re-enter the axon (Eq. (45)). If more mitochondria re-enter the axon (which corresponds to a greater value of *α*), the flux of newly synthesized mitochondria entering the first demand site in our model is reduced to keep the total flux of mitochondria entering the first demand site (newly synthesized plus returning, Fig. 2) the same.

The reduction of parameter *p*_*s*_, which characterizes the rate of mitochondria transition to the stationary state (Fig. 2), to 0.1, resulted in a reduction of the total length of mitochondria per unit length in the stationary state by approximately a factor of 4, which is due to fewer mitochondria transitioning to the stationary state (Fig. S1b). The increase of the value of *α* to 0.3 did not change the distributions of the total length of mitochondria per unit axonal length (Fig. S1b). This is because, according to our assumptions, the change of *α* does not affect the flux of mitochondria entering the most proximal demand site.

Anterograde and retrograde fluxes between the compartments are also independent of the values of *α* and *p*_*s*_, and are uniform along the axon length (Fig. S2a,b).

### 3.2 Mean age and age density distributions of mitochondria

In Fig. 3, we use line graphs to depict the mean age of mitochondria versus the demand site number, which characterizes the distance from the soma (Fig. 1). It should be noted, however, that per ref. [25], we simulate the demand sites as highly confined segments within the axon where stationary mitochondria are present. For *p*_*s*_ = 0.4 and *α* = 0 the mean age of mitochondria increases gradually from the most proximal to the most distal demand site. The mean age of anterograde mitochondria is the smallest, the mean age of stationary mitochondria is in the middle, and the mean age of retrograde mitochondria is the greatest. The mean age of stationary mitochondria in the most distal demand site is approximately 21 hours (Fig. 3a).

**Fig. 3.**
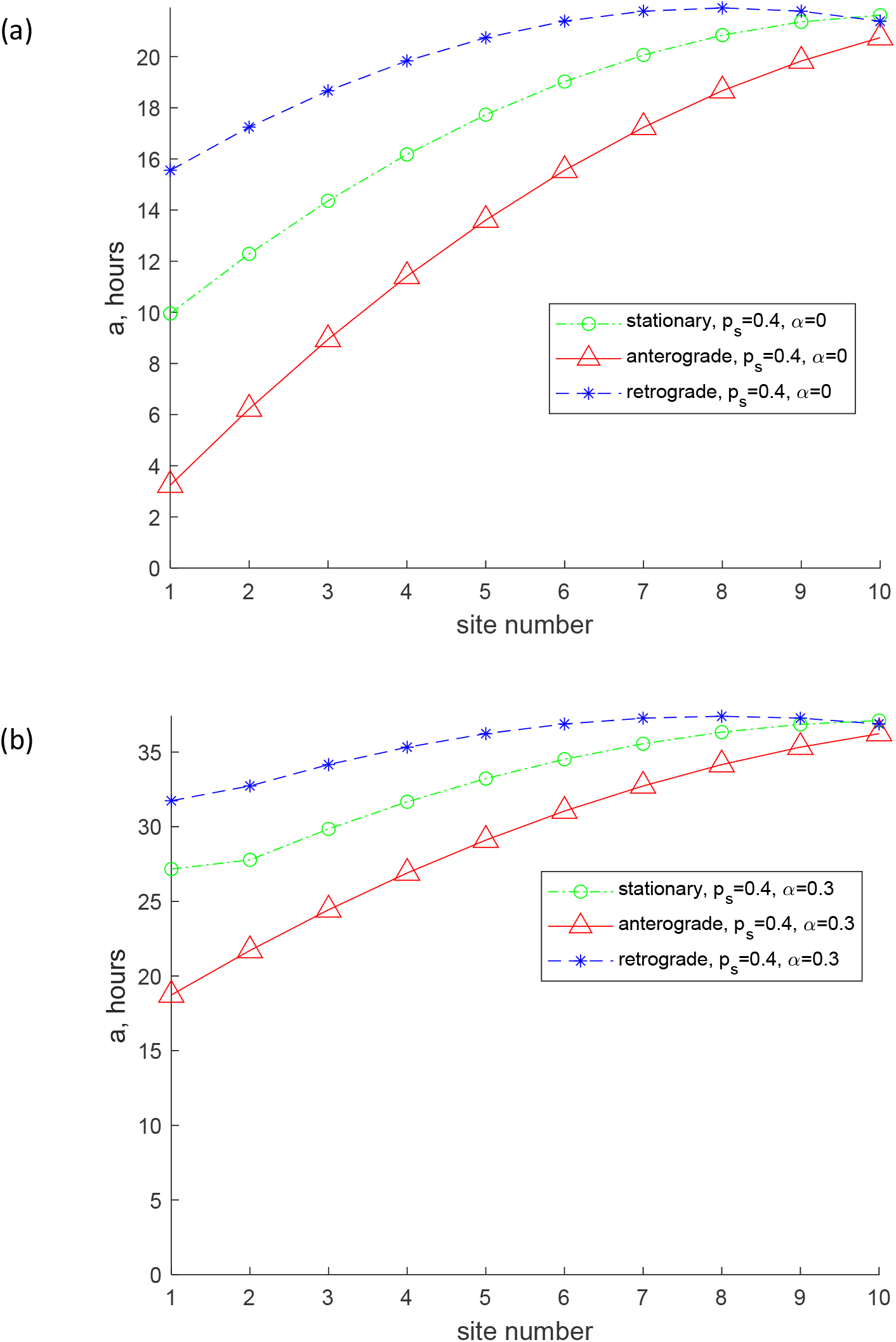
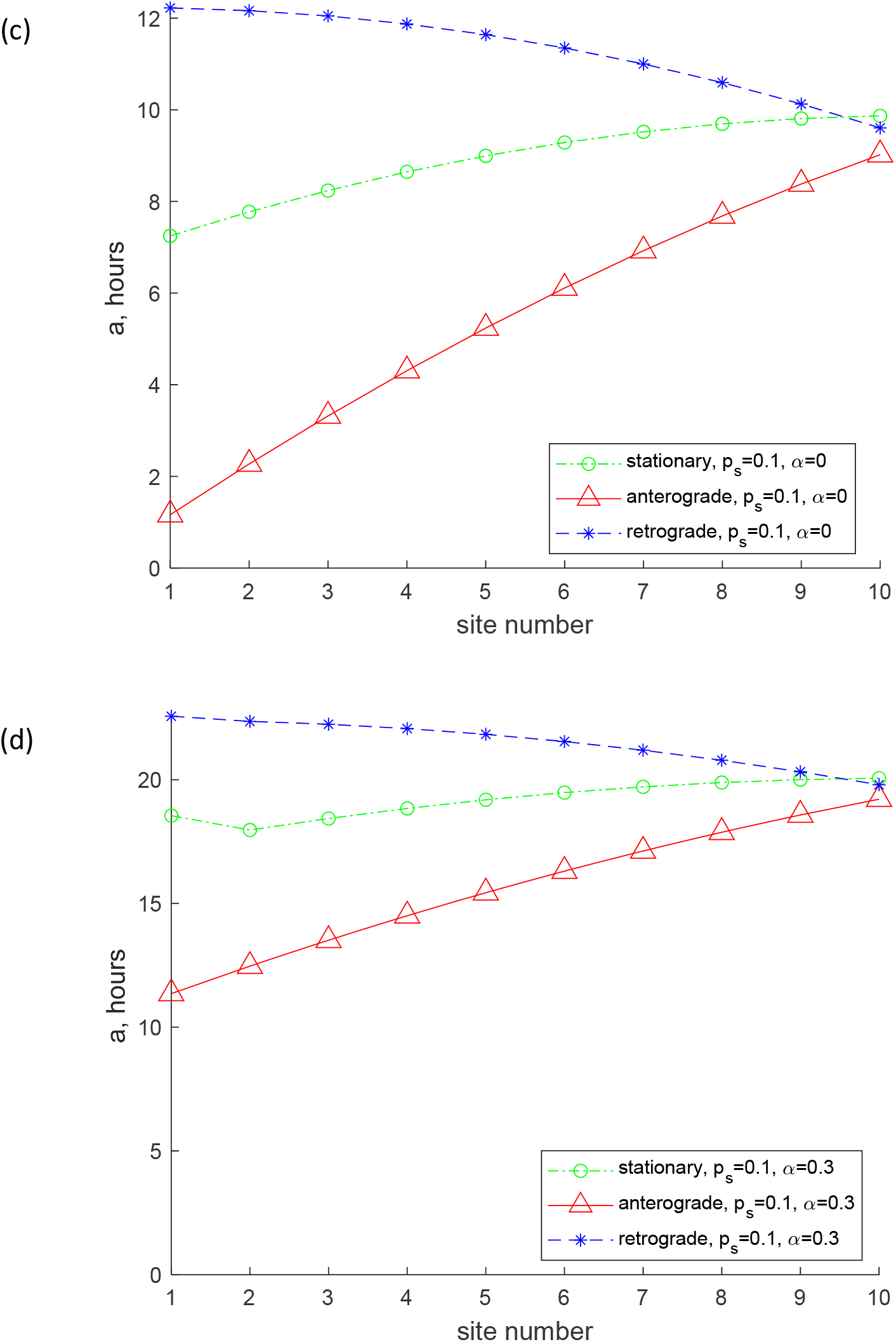
Mean age of stationary, anterogradely moving, and retrogradely moving mitochondria. (a) *p*_*s*_ = 0.4, *α* = 0 ; (b) *p*_*s*_ = 0.4, *α* = 0.3 ; (c) *p*_*s*_ = 0.1, *α* = 0 ; (d) *p*_*s*_ = 0.1, *α* = 0.3. Number of demand sites *N*=10.

Since the compartmental model does not consider the time it takes for mitochondria to transition between the compartments [42], it is necessary to prove that the age of mitochondria predicted by the model is less than the time it takes for mitochondria to travel from the soma to the axon tip without stops, *L*_*ax*_ / *v* :

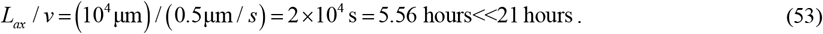

This means that a compartmental model can be used to simulate the transport of mitochondria in relatively short axons, as one simulated in this paper, of the length of 10^4^ μm.

The mean age for mitochondria for *p*_*s*_ = 0.4 and *α* = 0 roughly follows the same trend (an increase from the most proximal to the most distal demand site), but it is now much greater. The mean age of stationary mitochondria in the most distal demand site is now approximately 36 hours (Fig. 3b). This is because for *α* = 0.3 the flux of mitochondria entering the axon includes mitochondria that leave the axon moving retrogradely (Fig. 2). We thus successfully simulated the situation described in ref. [19] that suggests that older mitochondria appear in the axon due to return of mitochondria leaving the axon, which turn around and return to the axon in the anterograde component. It should be noted that the real situation is even more complicated, since mitochondria returning to the axon are most likely repaired in the soma by fusion with newly synthesized mitochondria. Furthermore, moving mitochondria can fuse with stationary mitochondria residing in the axon and exchange mitochondrial proteins [5,25,43]. This process will need to be simulated in future versions of the model.

For a smaller value of *p*_*s*_ (*p*_*s*_ = 0.1, *α* = 0, Fig. 3c), the mean age of mitochondria is significantly reduced. The mean age of stationary mitochondria in the most distal demand site is now approximately 10 h. This is because the probability of transitioning between moving states (anterograde or retrograde) and the stationary state (Fig. 2) is now much smaller. It is interesting that although the mean age of anterograde and stationary mitochondria increases from the most proximal to the most distal demand site, the age of retrograde mitochondria now decreases from the most proximal to the most distal demand site (Fig. 3c). This is because the exchange between moving and stationary mitochondria is now much less frequent (due to a smaller value of *p*_*s*_), and mitochondria mostly remain in the same kinetic state (anterograde, stationary, or retrograde). Retrograde mitochondria age as they return from the most distal demand site (site 10) to the most proximal demand site (site 1), which explains why the age of retrograde mitochondria is the largest in the most proximal demand site.

The increase of *α* (*p*_*s*_ = 0.1, *α* = 0.3, Fig. 3d) results in the increase of the mean age of mitochondria in all demand sites, but the trends stay the same. The mean age of the stationary mitochondria in the most distal demand site is now approximately 20 h. The increase in the mean age is due to the return of 30% of mitochondria exiting the most proximal demand site, which turn around and come back to the axon (Fig. 2).

The age density of mitochondria is important to characterize the ranges of mitochondria ages in different kinetic states (anterograde, stationary, and retrograde) in different demand sites. Fig. 4a compares the age densities of mitochondria computed for the base case (*p*_*s*_ = 0.4, *α* = 0, red lines) with age densities of mitochondria computed for the case of *p*_*s*_ = 0.4, *α* = 0.3, blue lines. The age densities of anterograde mitochondria are more narrow (Fig. 4a, see also Fig. 4b that shows a magnified view of Fig. 4a). Notably, the distribution of age densities widens for stationary mitochondria (Fig. 4c) and even wider for retrograde mitochondria (Fig. 4d). The shift of the age density curves towards older mitochondria, observed when *α* is increased from 0 to 0.3 (Fig. 4d), can be attributed to the reentry of previously aged retrograde mitochondria back into the axon. The presence of older mitochondria is consistent with the results reported in ref. [19]. The peaks on the curves displaying the age density distributions are shifted toward older mitochondria for more distal demand sites (Fig. 4).

**Fig. 4.**
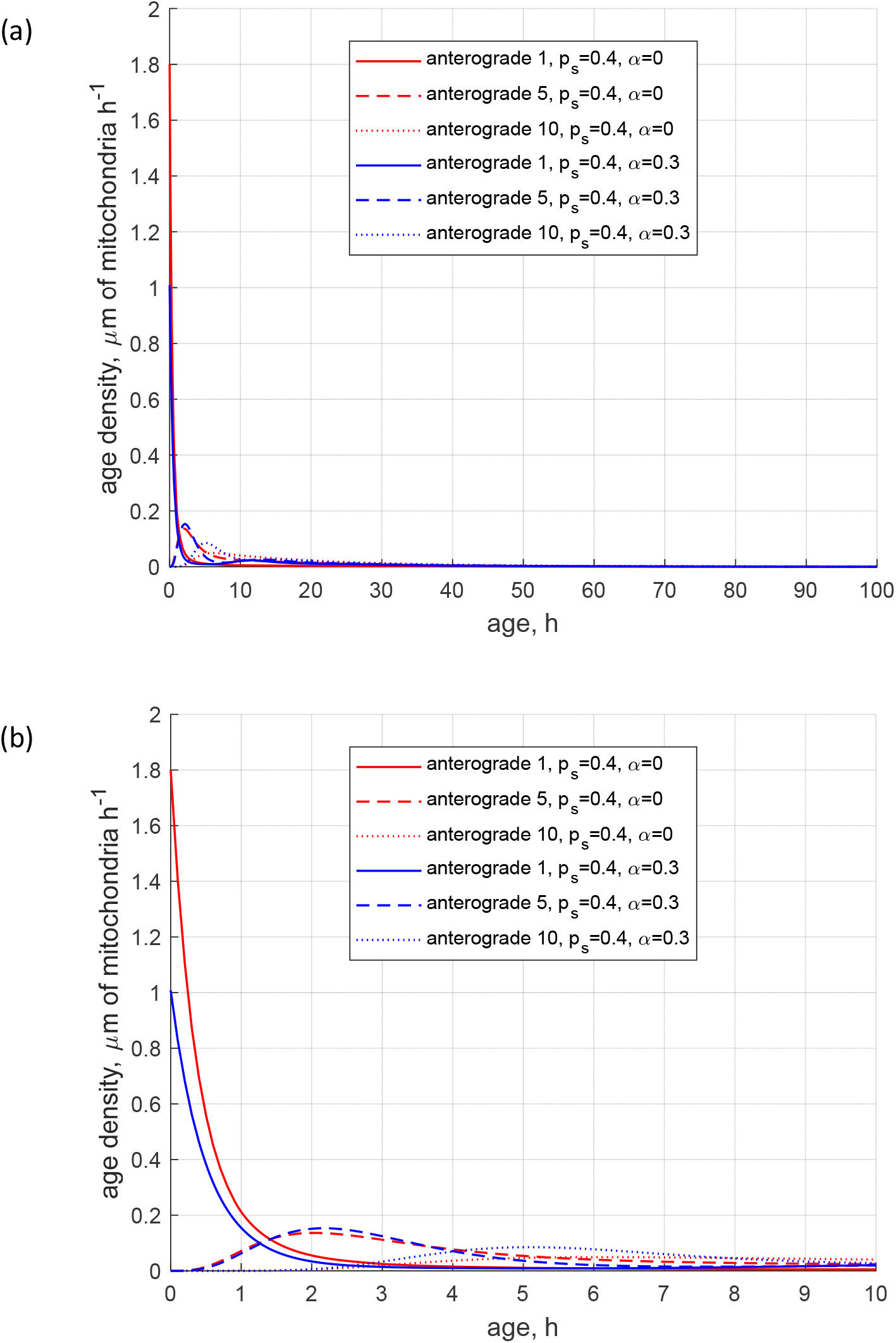
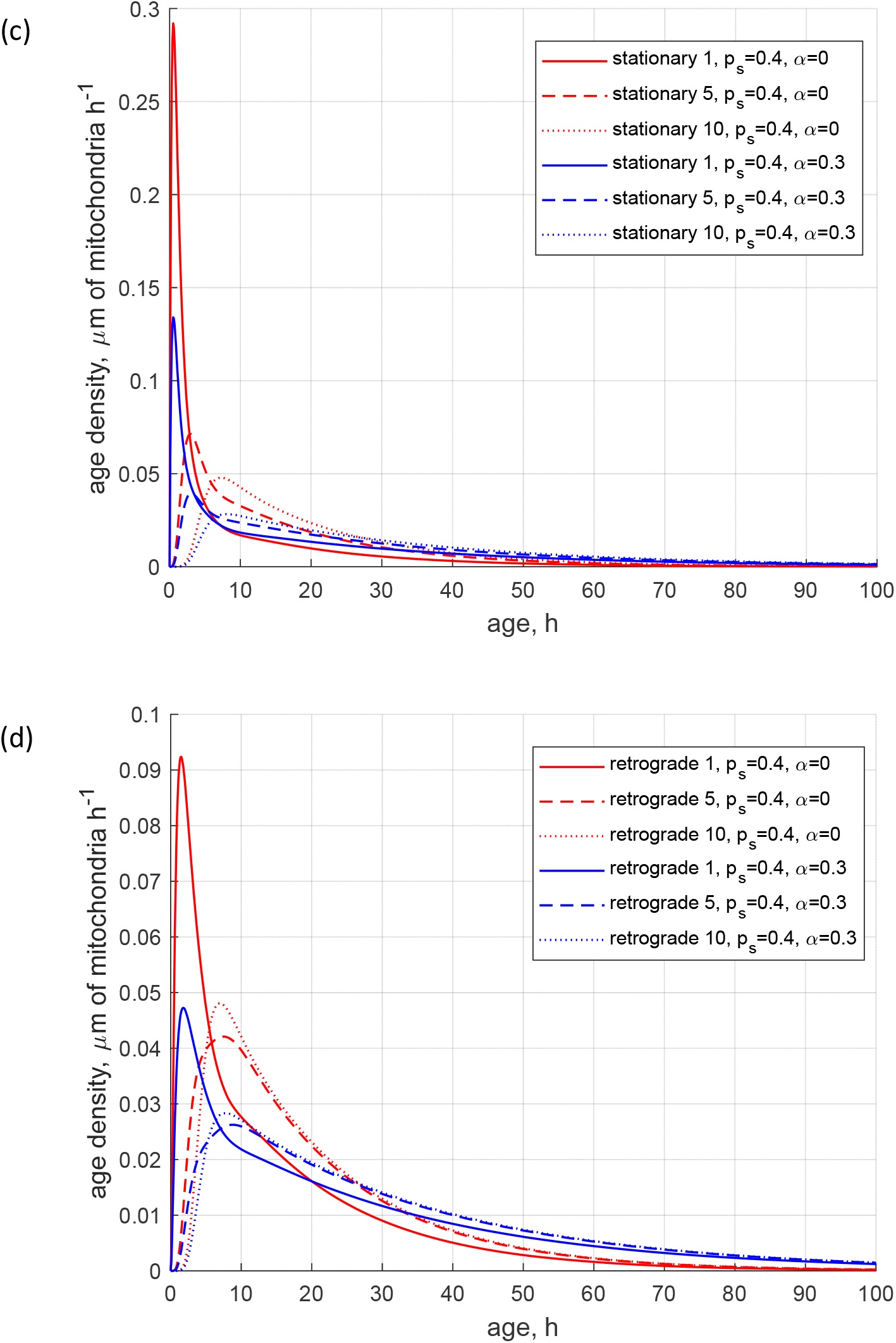
(a) Age density of anterogradely moving mitochondria in various demand sites. (b) Similar to Fig. 4a, this graph magnifies a specific age range of [0 10 hours] of the *x*-axis. (c) Age density of mitochondria in the stationary state in various demand sites. (d) Age density of retrogradely moving mitochondria in various demand sites. Two sets of parameter values: *p*_*s*_ = 0.4, *α* = 0 (base case) and *p*_*s*_ = 0.4, *α* = 0.3. Number of demand sites *N*=10.

The reduction of the rate of mitochondria transition to the stationary state (characterized by *p*_*s*_) makes the age density distribution of mitochondria (*p*_*s*_ = 0.1, *α* = 0, blue curves, Fig. 5) quite complex. The most noticeable change is the bimodal distribution that is visible in Fig. 5d in the solid blue curve displaying the age density of retrograde mitochondria in the most proximal demand site (site 1). The first peak occurs at approximately 1 h. It is explained by mitochondria that recently entered the axon, but then transitioned to the stationary state, and then to the retrograde state (Fig. 2). The second peak occurs at approximately 11 h. It is explained by older mitochondria that travelled to the axon tip and then returned to the most proximal site.

**Fig. 5.**
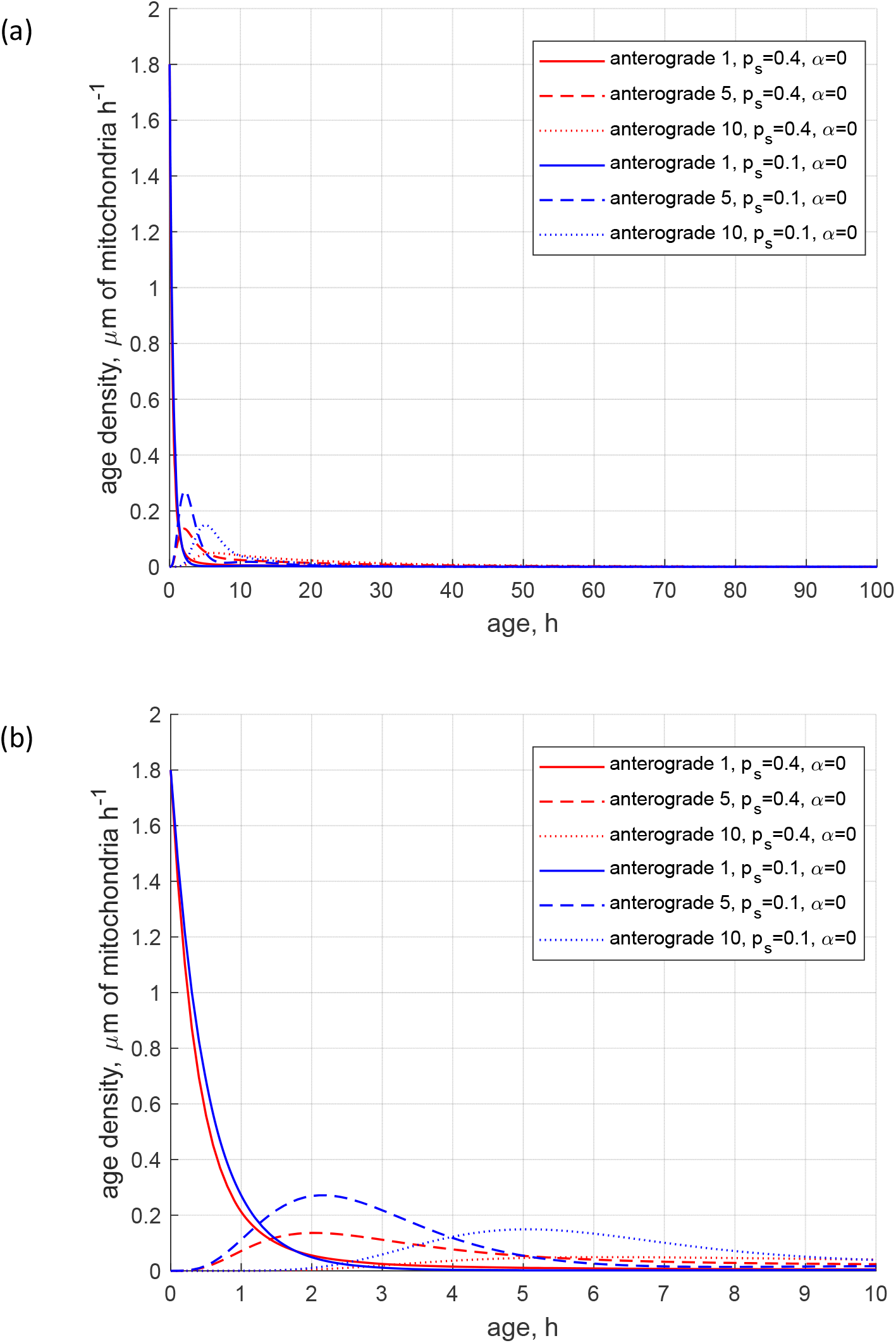
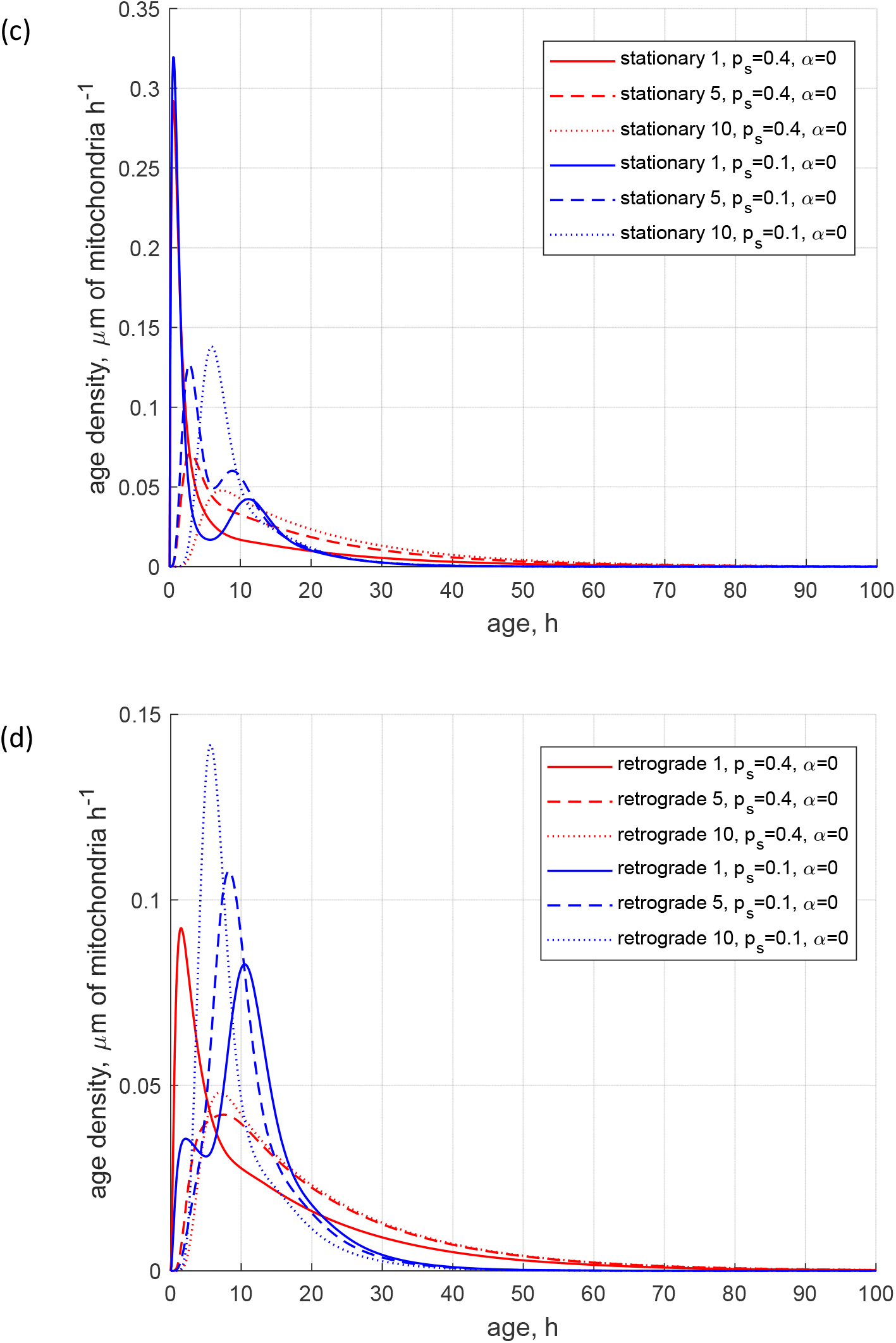
(a) Age density of anterogradely moving mitochondria in various demand sites. (b) Similar to Fig. 5a, this graph magnifies a specific age range of [0 10 hours] of the *x*-axis. (c) Age density of mitochondria in the stationary state in various demand sites. (d) Age density of retrogradely moving mitochondria in various demand sites. Two sets of parameter values: *p*_*s*_ = 0.4, *α* = 0 (base case) and *p*_*s*_ = 0.1, *α* = 0. Number of demand sites *N*=10.

The increase of the value of *α* (more mitochondria return to recirculation in the axon) while keeping *p*_*s*_ low (small transition rate to the stationary state) makes the tails in age density distributions longer, which means the presence of older mitochondria in the axon, see ref. [35] (*p*_*s*_ = 0.1, *α* = 0.3, blue curves in Fig. S3d). Interestingly, the first peak on the solid blue curve in Fig. S3d, displaying the age density of retrograde mitochondria in site 1 (at approximately 1 h), has almost disappeared; it degraded nearly to the horizontal slope of the curve (compare Fig. S3d with Fig. 5d). All age density distributions in Figs. 4, 5, S3, and in particular in Fig. S3d, are skewed right. This indicates that although the mean ages of mitochondria are of the order of 20 hours (see Fig. 3d, which shows mean ages of mitochondria that correspond to the age density distributions depicted by blue lines in Fig. S3d), there are also much older mitochondria present in the axon, which stay in the axon several times longer than 20 h, probably completing several circulations or staying a long time in the stationary state before returning to the mobile pool.

We have confirmed that the bimodal age density distribution of retrograde mitochondria is not exclusive to 10 demand sites. In Figs. S4 and S5 in Supplemental Materials, we analyzed the age density distributions of mitochondria in an axon consisting of five demand sites (*N*=5). The bimodal distributions observed for retrograde mitochondria in the most proximal demand site are visible in Figs. S4d and S5d. Our hypothesis suggests that the bimodal distribution arises from the presence of retrograde mitochondria with two distinct ages: (i) relatively young mitochondria that stop and become retrograde, and (ii) relatively old mitochondria that travel to the end of the axon, reverse their direction, and return. The difference in ages between these two types of mitochondria is most prominent in the most proximal demand site, resulting in the occurrence of a bimodal distribution. In more distant demand sites, the bimodal pattern may become less pronounced.

### 3.3 Sensitivity of the mean age of mitochondria in demand sites to *p*_*s*_ and *α*

This investigation was performed by computing local sensitivity coefficients, which are first-order partial derivatives of the observables with respect to model parameters [44-47]. For example, the sensitivity coefficient of the mean age of resident mitochondria in demand sites to parameter *α* at steady-state (ss) was calculated as follows:

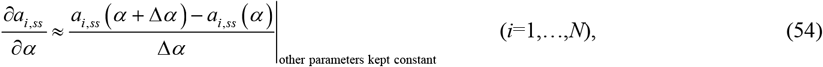

where Δ*α* =10^−1^*α*(we tested the accuracy by using various step sizes).

To make sensitivity coefficients independent of the magnitude of the parameter whose sensitivity was tested, we calculated non-dimensional relative sensitivity coefficients [45,48], which were defined as (for example):

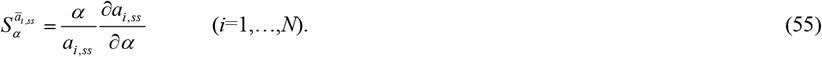

The dimensionless sensitivity of the mean age of mitochondria to the probability that mobile mitochondria would transition to a stationary state at a demand site, *p*_*s*_, is positive in all demand sites (Fig. 6a). This is because mitochondria spend more time in the stationary state when the stopping probability is larger.

**Fig. 6.**
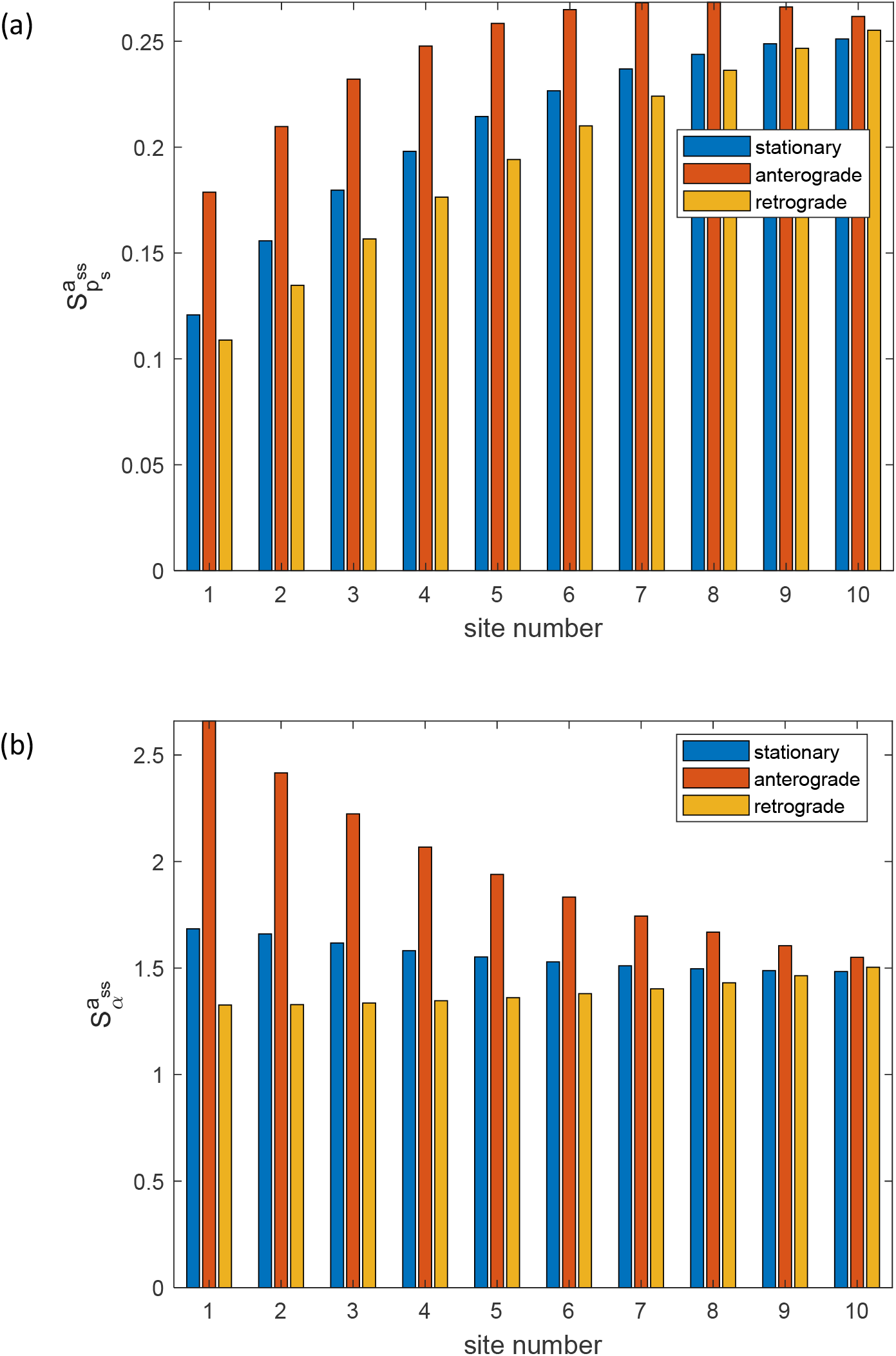
(a) Dimensionless sensitivity of the mean age of mitochondria to the probability that mobile mitochondria would transition to a stationary state at a demand site, *p*_*s*_, versus the site number. Computations were performed with Δ*p*_*s*_ = 10^−1^ *p*_*s*_. A close result was obtained for Δ*p*_*s*_ = 10^−2^ *p*_*s*_. (b) Dimensionless sensitivity of the mean age of mitochondria to the portion of mitochondria that return to the axon after exiting the axon, *α*, versus the site number. Computations were performed with Δ*α* = 10^−1^*α*. A close result was obtained for Δ*α* = 10^−2^*α*. Sensitivities are analyzed around *p*_*s*_ = 0.1 s^-1^, *α* = 0.3. Number of demand sites *N*=10.

The dimensionless sensitivity of the mean age of mitochondria to the portion of mitochondria that return to the axon after exiting the axon, *α*, is also positive in all demand sites (Fig. 6b). This is because older mitochondria, which already circulated in the axon, re-enter the axon when the value of *α* is larger than zero. The number of older mitochondria in the axon thus increases. This is consistent with the results depicted in Fig. 3 (compare the results in Fig. 3a and Fig. 3b, and also the results in Fig. 3c and Fig. 3d). It is interesting to note that the mean age of stationary mitochondria is approximately 10 times more sensitive to *α* than to *p*_*s*_ (compare Fig. 6b with Fig. 6a). We examined whether this conclusion remains consistent regardless of the number of demand sites. The findings depicted in Fig. S6, computed for five demand sites (*N*=5), reveal that the sensitivity to *α* is independent of the number of compartments, while the sensitivity to *p*_*s*_ scales linearly with the number of compartments in the axon. This outcome arises from the fact that *p*_*s*_ represents the stopping probability in each compartment. Notably, this result implies that for a very long axon with many compartments, the age of mitochondria may be highly sensitive to the stopping probability.

## 4. Discussion, limitations of the model, and future directions

We developed a model that accounts for the return of a portion of mitochondria (characterized by parameter *α*) back to the axon after mitochondria complete circulation in the axon. We investigated how the mean age and age density distribution of mitochondria depend on *α*. We also investigated the dependence of the same quantities on *p*_*s*_.

Bimodal age density distributions are found for a smaller value of *p*_*s*_ (*p*_*s*_ = 0.1). The peak corresponding to older mitochondria on the curve displaying the age density distribution of retrograde mitochondria is explained by mitochondria that traveled to the tip of the axon and then traveled back in the retrograde component. The peak corresponding to younger mitochondria on the curve displaying the age density distribution of retrograde mitochondria is explained by mitochondria that transitioned from the anterograde state to the state occupied by stationary mitochondria and then to the retrograde state.

We found that the mean age of stationary mitochondria is very sensitive to parameter *α*. It is an order of magnitude less sensitive to parameter *p*_*s*_. We also investigated how the sensitivity of the mean age of stationary mitochondria depends on the number of compartments in the axon. The sensitivity to the stopping probability, *p*_*s*_, increases proportionally with the number of compartments in the axon. This suggests that in an extremely long axon with a large number of compartments, the age of mitochondria could be highly influenced by the stopping probability.

The prediction of a broader age density distribution for retrograde mitochondria compared to anterograde mitochondria could in principle be accessible with experimental measurements.

Future research should aim to explore the feasibility of analytically solving equations of our model to obtain explicit expressions for the age densities of mitochondria in anterograde, stationary, and retrograde kinetic states. Future modeling work should prove that the time-scale for transitioning between compartments in our model is comparable to the time-scale for mixing within an individual compartment. To investigate the time-scale for mixing within an individual compartment, it may be necessary to develop a model that would treat mitochondria as individual particles [25,49]. The utilization of a discrete model would provide much more detailed information on mitochondria trajectories and age. A promising extension of the current approach would also involve incorporating the stochastic nature of mitochondrial transport. Furthermore, in future research, a model of fusion/fission of mitochondria should be developed. This is especially important because fusion with newly synthesized mitochondria contributes to the replenishment of older mitochondria with new mitochondrial proteins [19]. Mitochondria removal via mitophagy [50,51] also needs to be incorporated in future models. Some mitochondria degradation may occur in axons locally [52,53]. The effect of this possibility should be incorporated into future models. Future research should also investigate the effect of domain discretization (using a different number of compartments) on the predictions of the developed model.

## Acknowledgment

IAK acknowledges the fellowship support of the Paul and Daisy Soros Fellowship for New Americans and the NIH/National Institute of Mental Health (NIMH) Ruth L. Kirchstein NRSA (F30 MH122076-01). AVK acknowledges the support of the National Science Foundation (award CBET-2042834) and the Alexander von Humboldt Foundation through the Humboldt Research Award.

## Ethical Statement

None.

## Supplemental Materials

### S1. Alternative form of governing equations

An alternative way of stating the conservation equations for the total length of mitochondria in the axon, developed in ref. [28], is as follows. The solution of these alternative governing equations is identical to the solution of vector Eq. (1).

Stating the conservation of the total length of stationary mitochondria in the most proximal demand site (site 1, Fig. 2) gives the following equation:

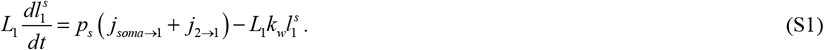

Stating the conservation of the total length of anterograde mitochondria in demand site 1 gives the following equation (Fig. 2):

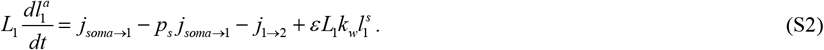

Stating the conservation of the total length of retrograde mitochondria in demand site 1 results in the following equation (Fig. 2):

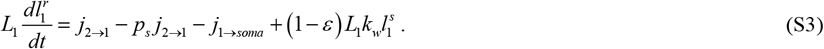

Stating the conservation of the total length of stationary mitochondria in the *i*th demand site (*i*=2,…,*N*-1) leads to the following equation (Fig. 2):

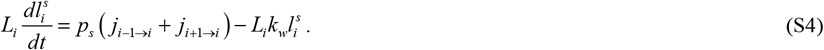

Stating the conservation of the total length of anterograde mitochondria in the *i*th demand site (*i*=2,…,*N*-1) leads to the following equation (Fig. 2):

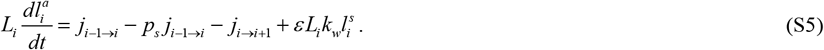

Stating the conservation of the total length of retrograde mitochondria in the *i*th demand site (*i*=2,…,*N*-1) results in the following equation (Fig. 2):

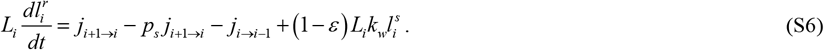

As suggested in ref. [25], mitochondria that reach the distal end of the axon instantaneously switch from anterograde motors (kinesins) to retrograde motors (dyneins) and continue moving retrogradely in the axon. This is simulated by *j*_*N* ⟶3*N*_ flux (Fig. 2). Stating the conservation of the total length of stationary mitochondria in the most distal demand site (site *N*) leads to the following equation (Fig. 2):

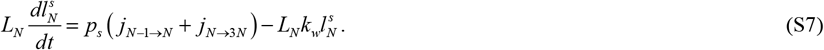

Stating the conservation of the total length of anterograde mitochondria in the most distal demand site (site *N*) leads to the following equation (Fig. 2):

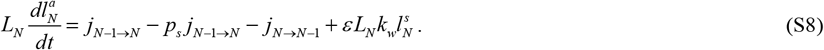

Stating the conservation of the total length of retrograde mitochondria in the most distal demand site (site *N*) leads to the following equation (Fig. 2):

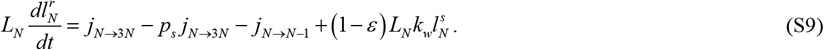

To obtain steady-state solutions, we set the left-hand side of Eqs. (S1)-(S9) to zero and solved the obtained system of linear equations for the total length of mitochondria per unit length of the axon in each compartment. Steady-state solutions of Eqs. (S1)-(S9) coincide with those displayed in Fig. S1, which validates our implementation of matrix B given by Eqs. (4)-(34) and Eq. (47).

### S2. Supplementary tables

**Table S1.**
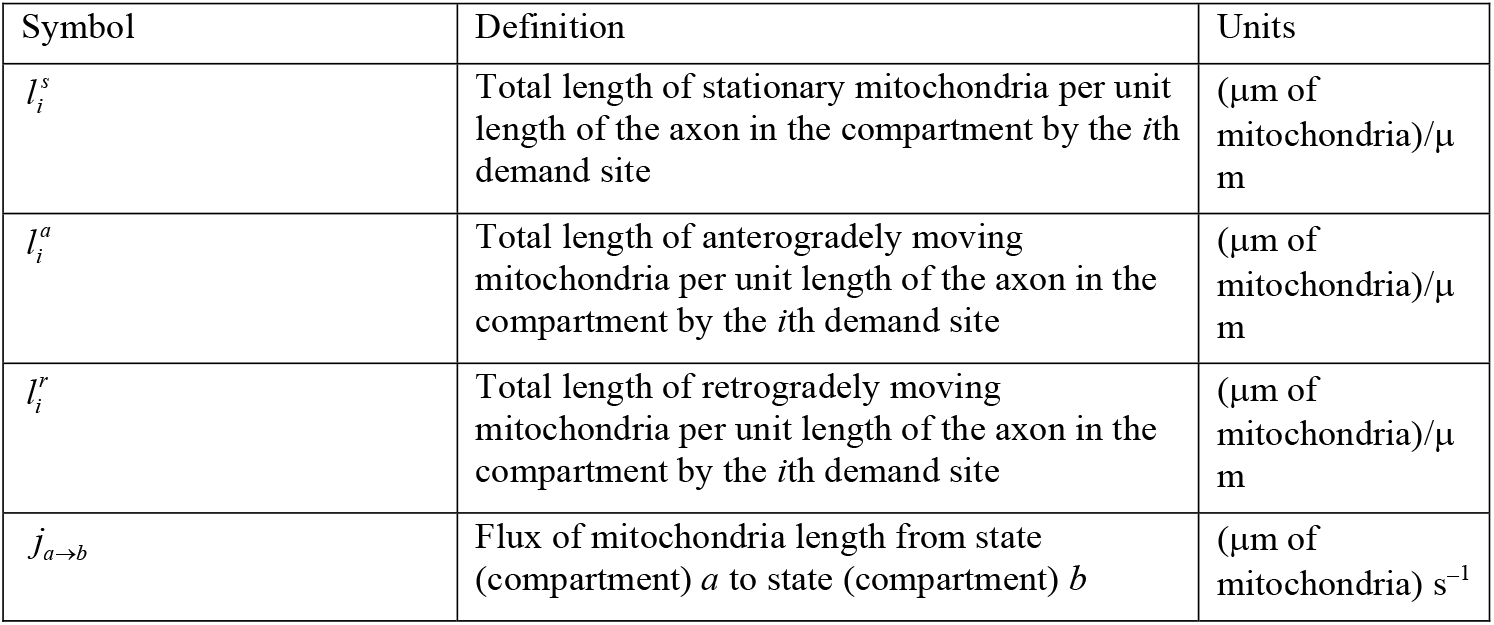
Dependent variables in the model of mitochondrial transport and accumulation in the axon.

| Symbol | Definition | Units |
| --- | --- | --- |
| $l_i^s$ | Total length of stationary mitochondria per unit length of the axon in the compartment by the $i$th demand site | ($\mu\text{m}$ of mitochondria)/$\mu\text{m}$ |
| $l_i^a$ | Total length of anterogradely moving mitochondria per unit length of the axon in the compartment by the $i$th demand site | ($\mu\text{m}$ of mitochondria)/$\mu\text{m}$ |
| $l_i^r$ | Total length of retrogradely moving mitochondria per unit length of the axon in the compartment by the $i$th demand site | ($\mu\text{m}$ of mitochondria)/$\mu\text{m}$ |
| $j_{a \rightarrow b}$ | Flux of mitochondria length from state (compartment) $a$ to state (compartment) $b$ | ($\mu\text{m}$ of mitochondria) $\text{s}^{-1}$ |

**Table S2.**
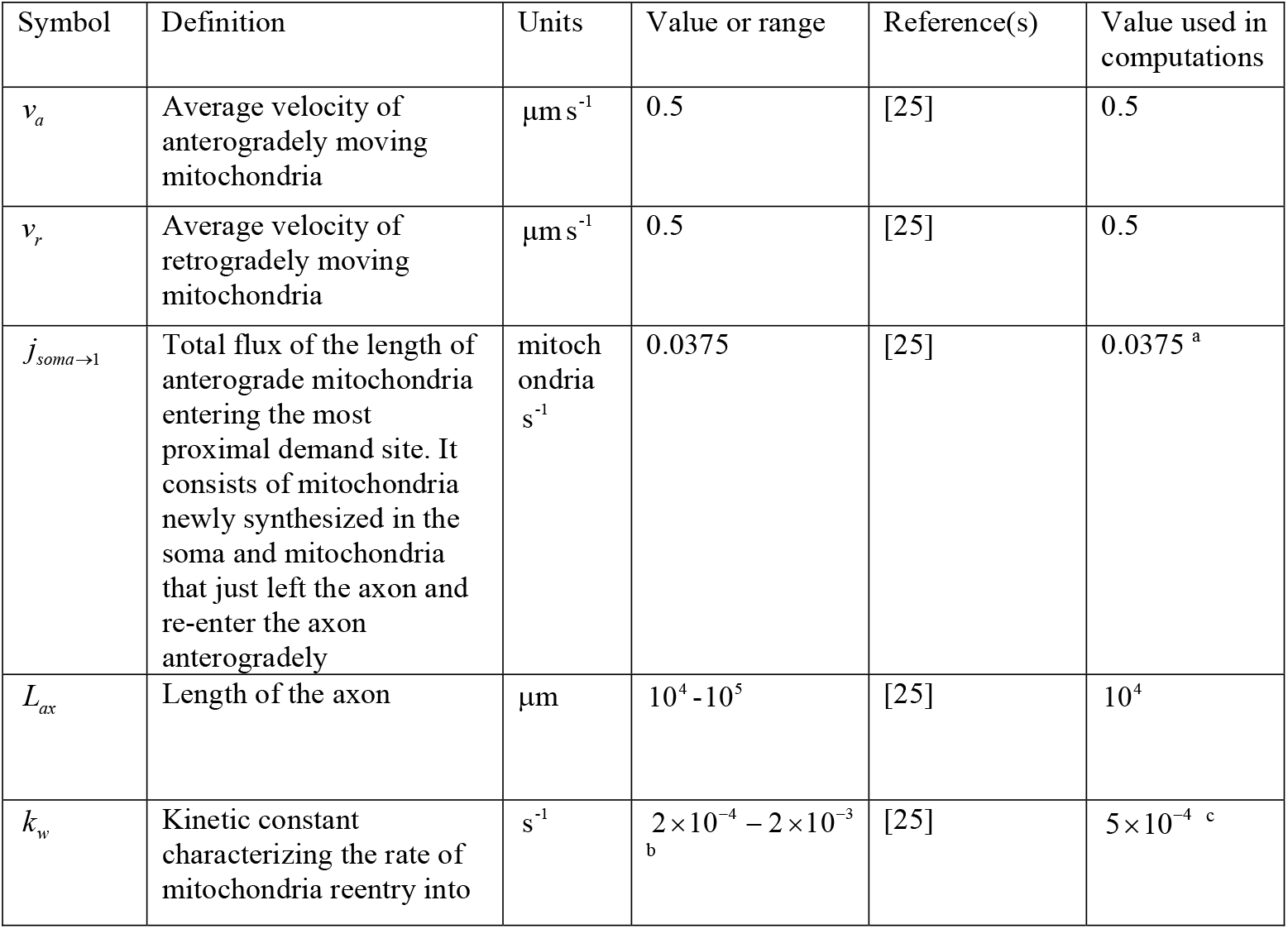
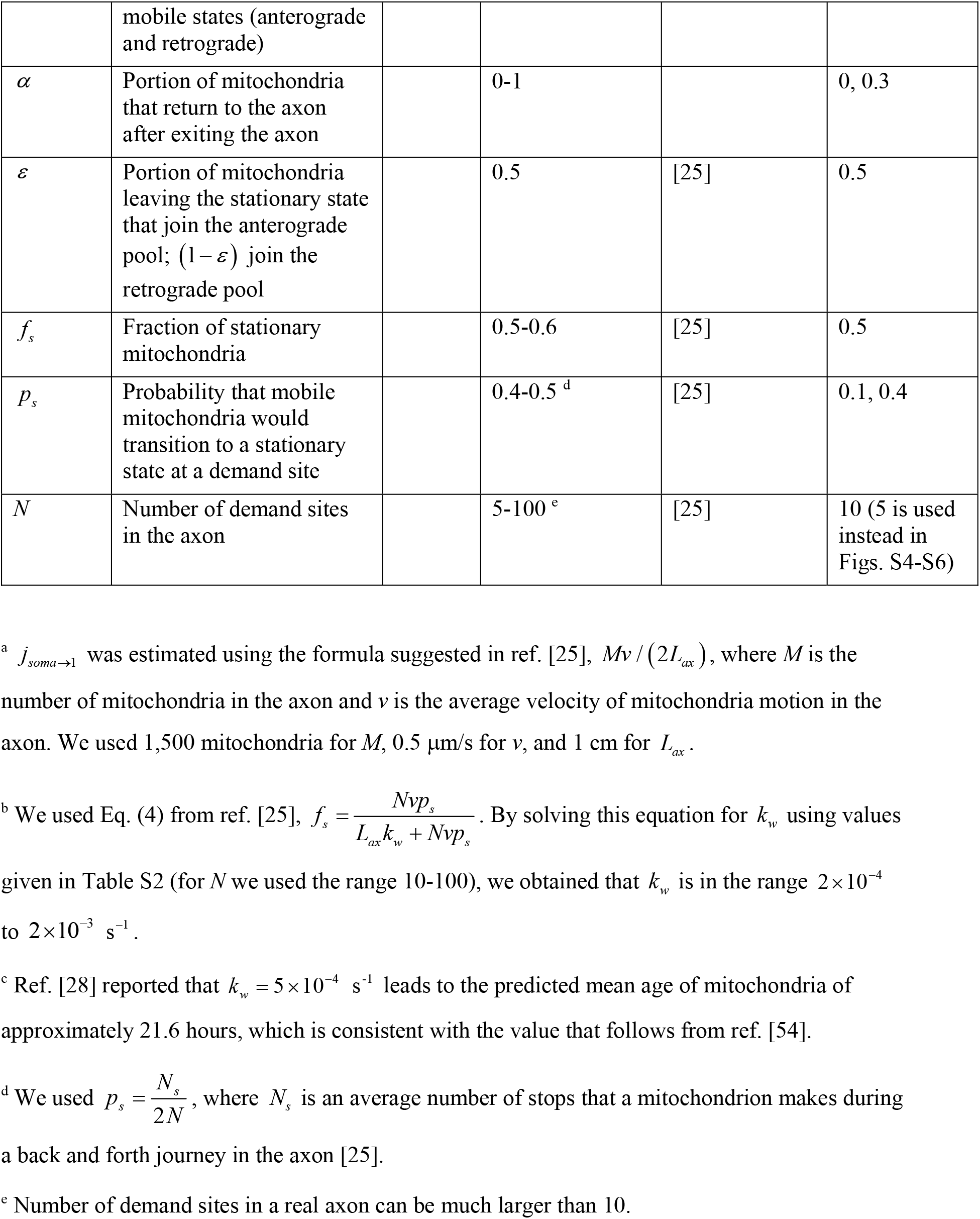
Parameters characterizing mitochondrial transport and accumulation in the axon.

### S3. Supplementary figures

**Fig. S1.**
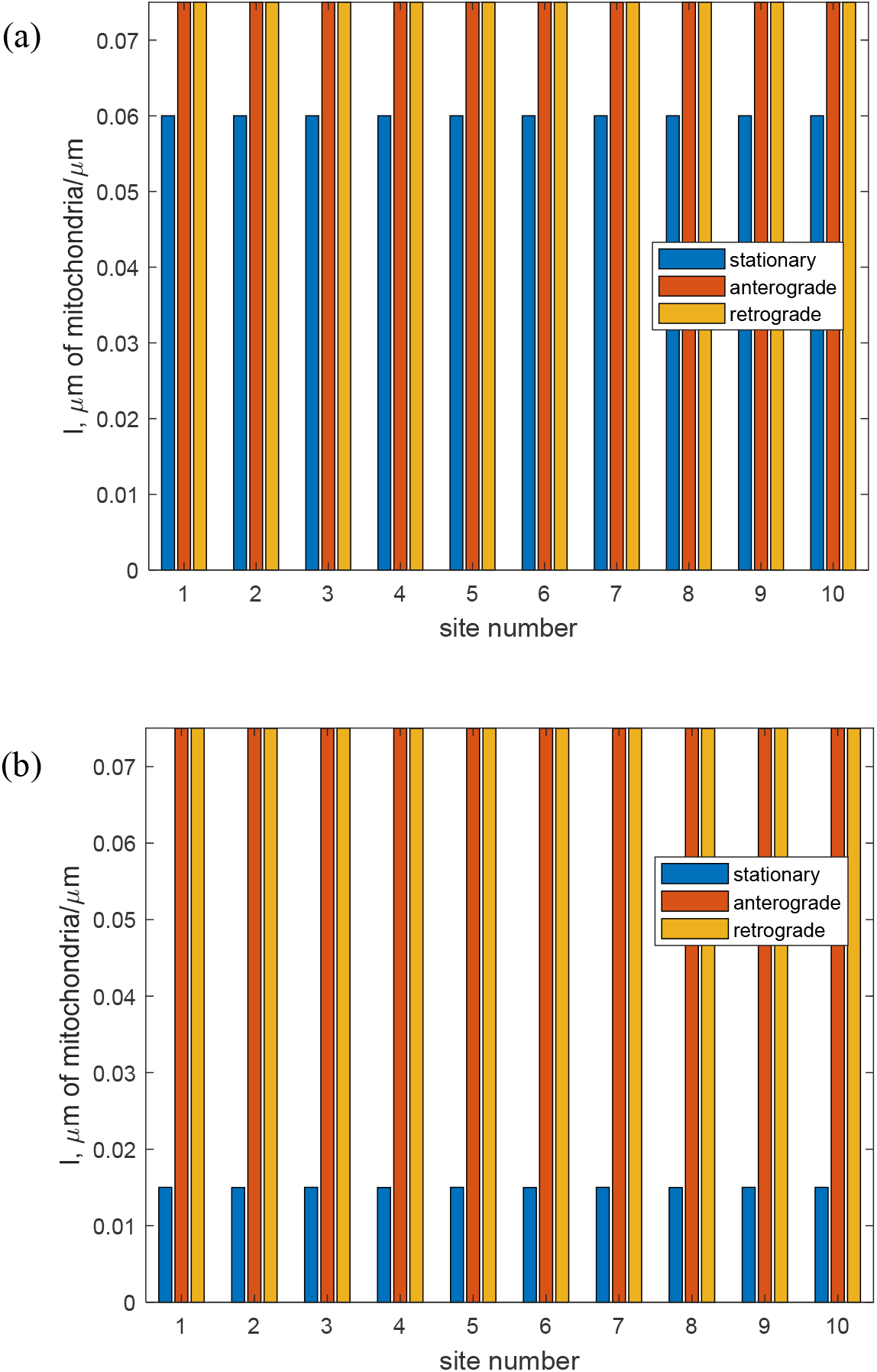
Steady-state values of the total length of stationary, anterogradely moving, and retrogradely moving mitochondria per unit length of the axon in the compartment by the *i*th demand site. (a) *p*_*s*_ = 0.4, the results are independent of the value of *α* ; (b) *p*_*s*_ = 0.1, the results are independent of the value of *α*. Number of demand sites *N*=10.

**Fig. S2.**
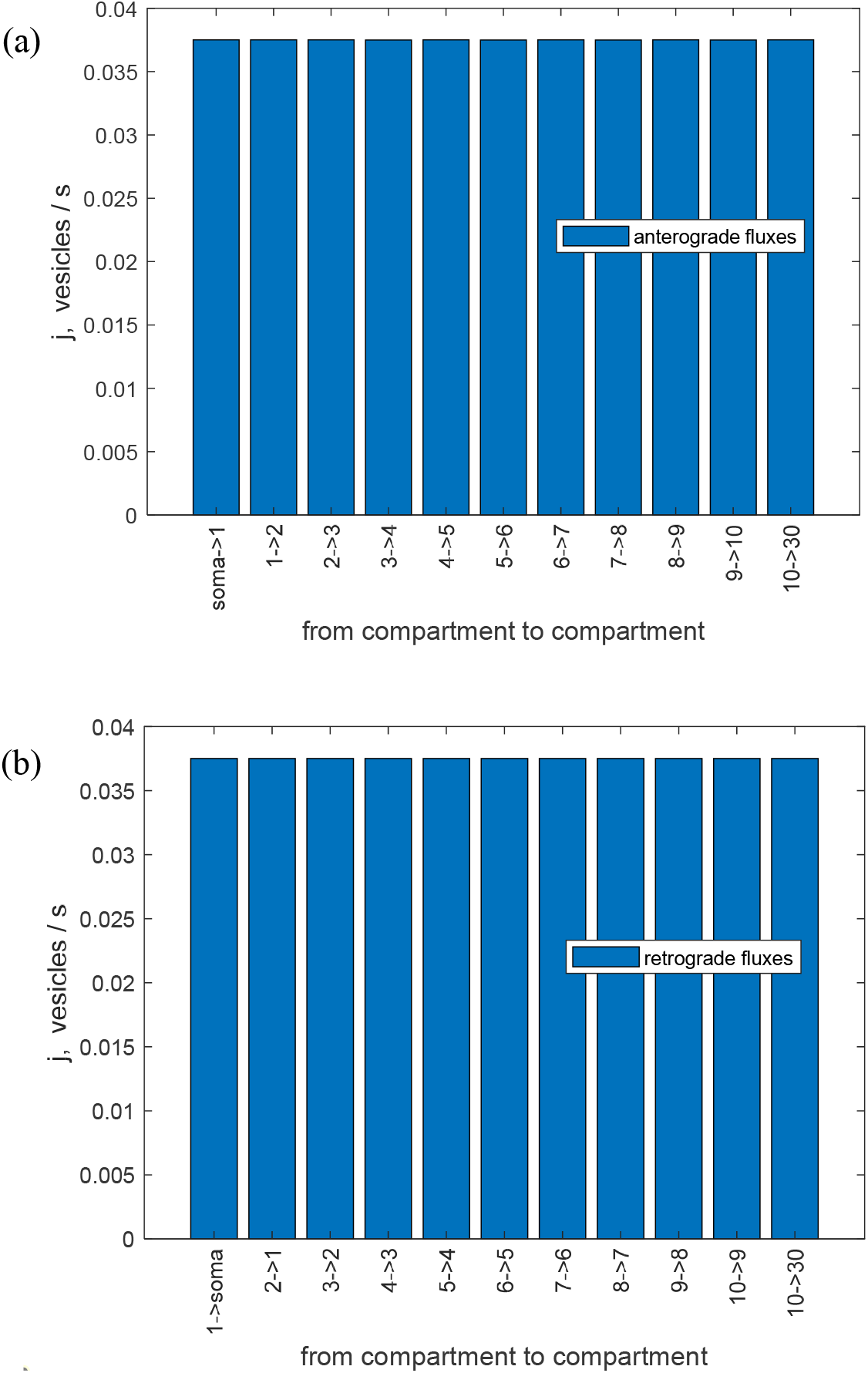
(a) Anterograde and (b) retrograde fluxes of mitochondria traveling between the compartments. The results are independent of the values of *p*_*s*_ and *α*. Number of demand sites *N*=10.

**Fig. S3.**
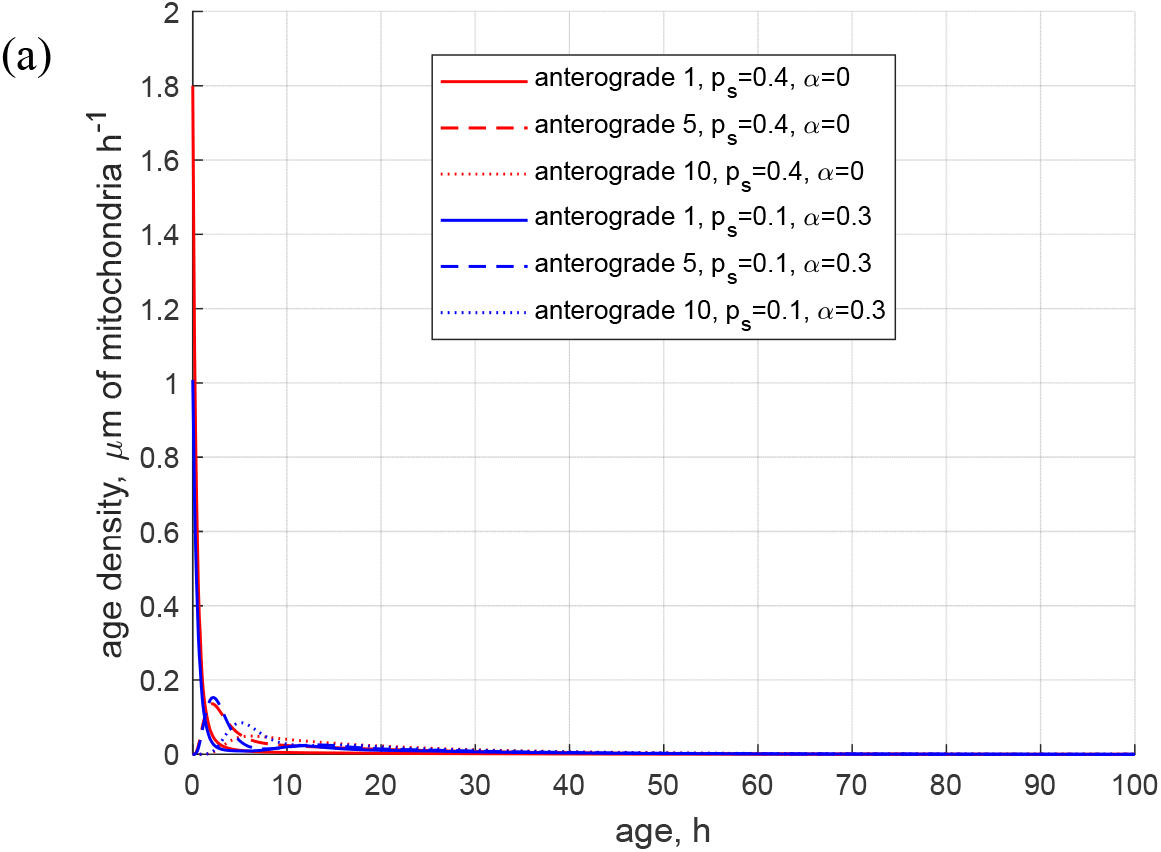
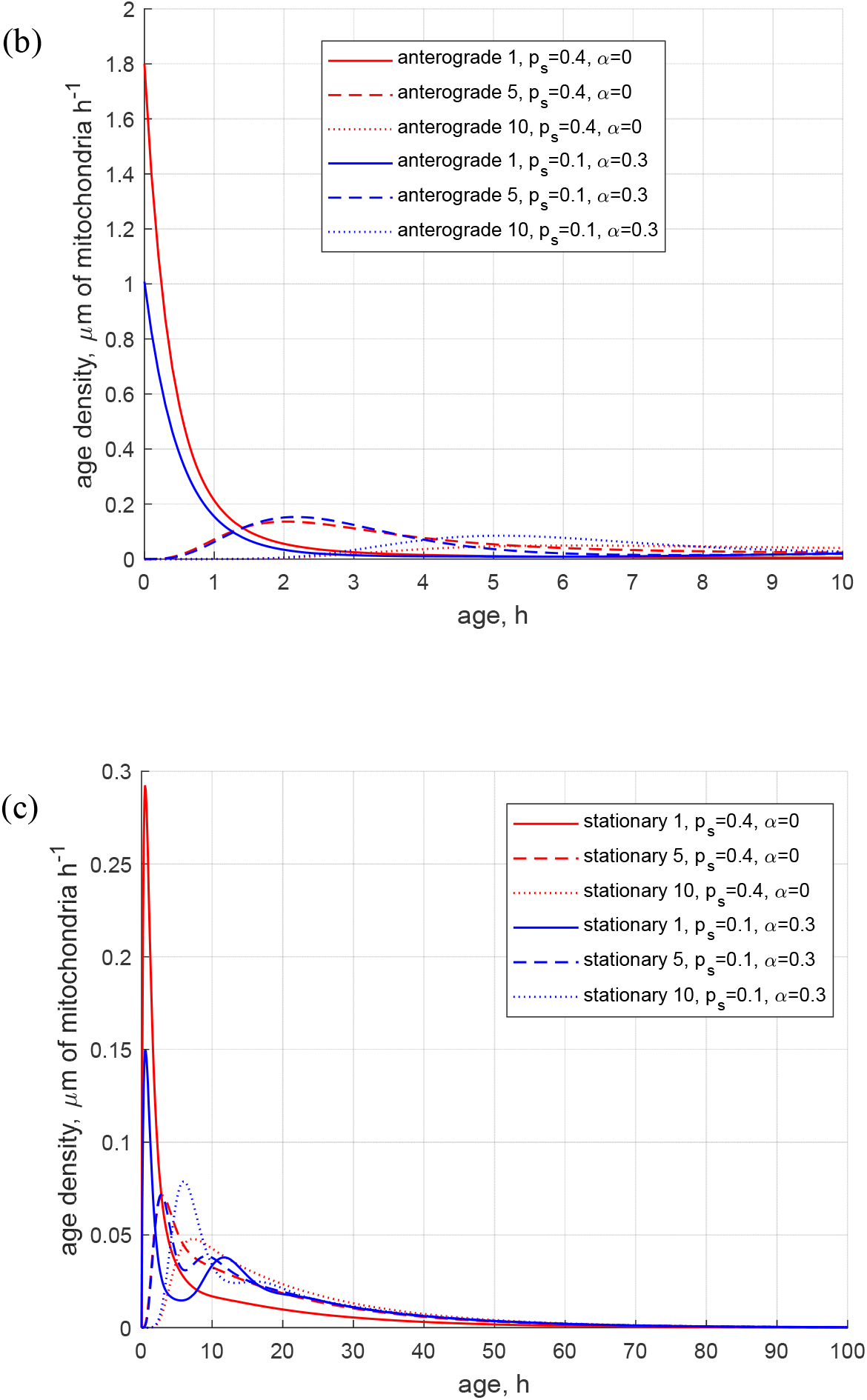
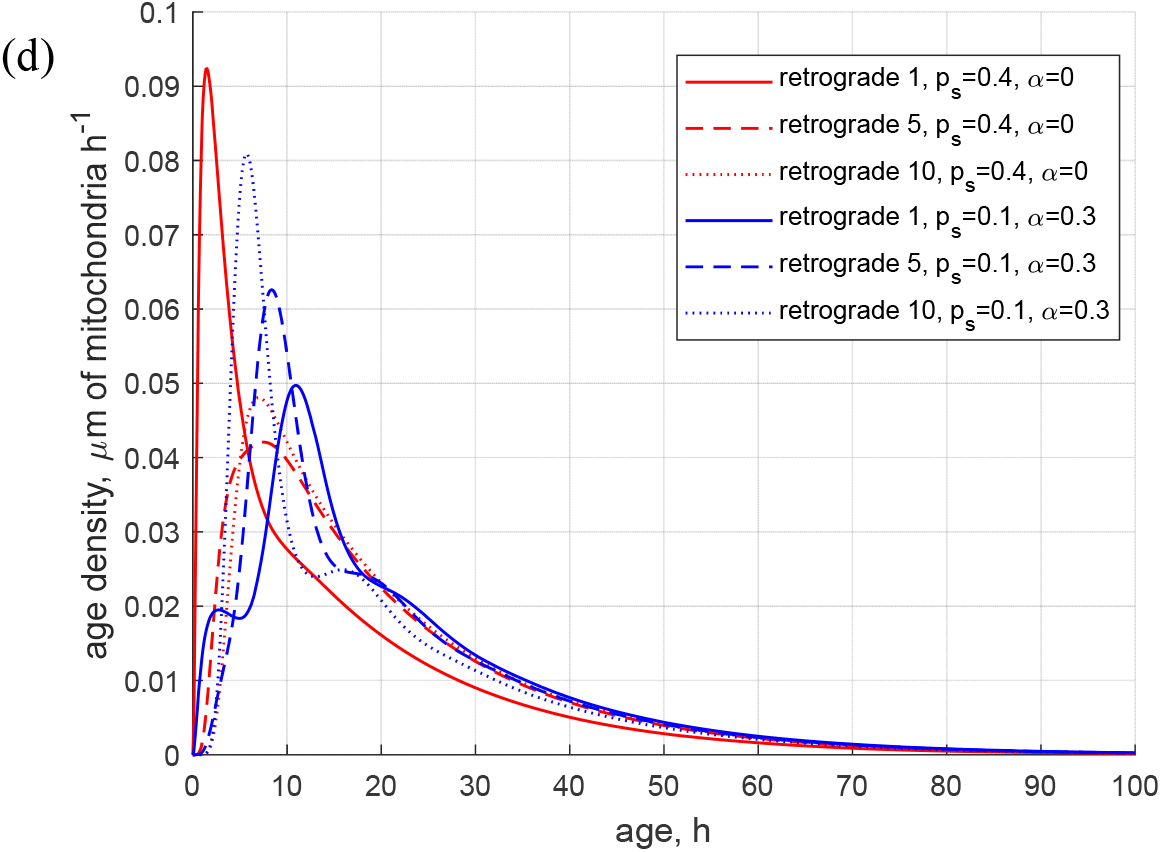
(a) Age density of anterogradely moving mitochondria in various demand sites. (b) Similar to Fig. S3a, this graph magnifies a specific age range of [0 10 hours] of the *x*-axis. (c) Age density of mitochondria in the stationary state in various demand sites. (d) Age density of retrogradely moving mitochondria in various demand sites. Two sets of parameter values: *p*_*s*_ = 0.4, *α* = 0 (base case) and *p*_*s*_ = 0.1, *α* = 0.3. Number of demand sites *N*=10.

**Fig. S4.**
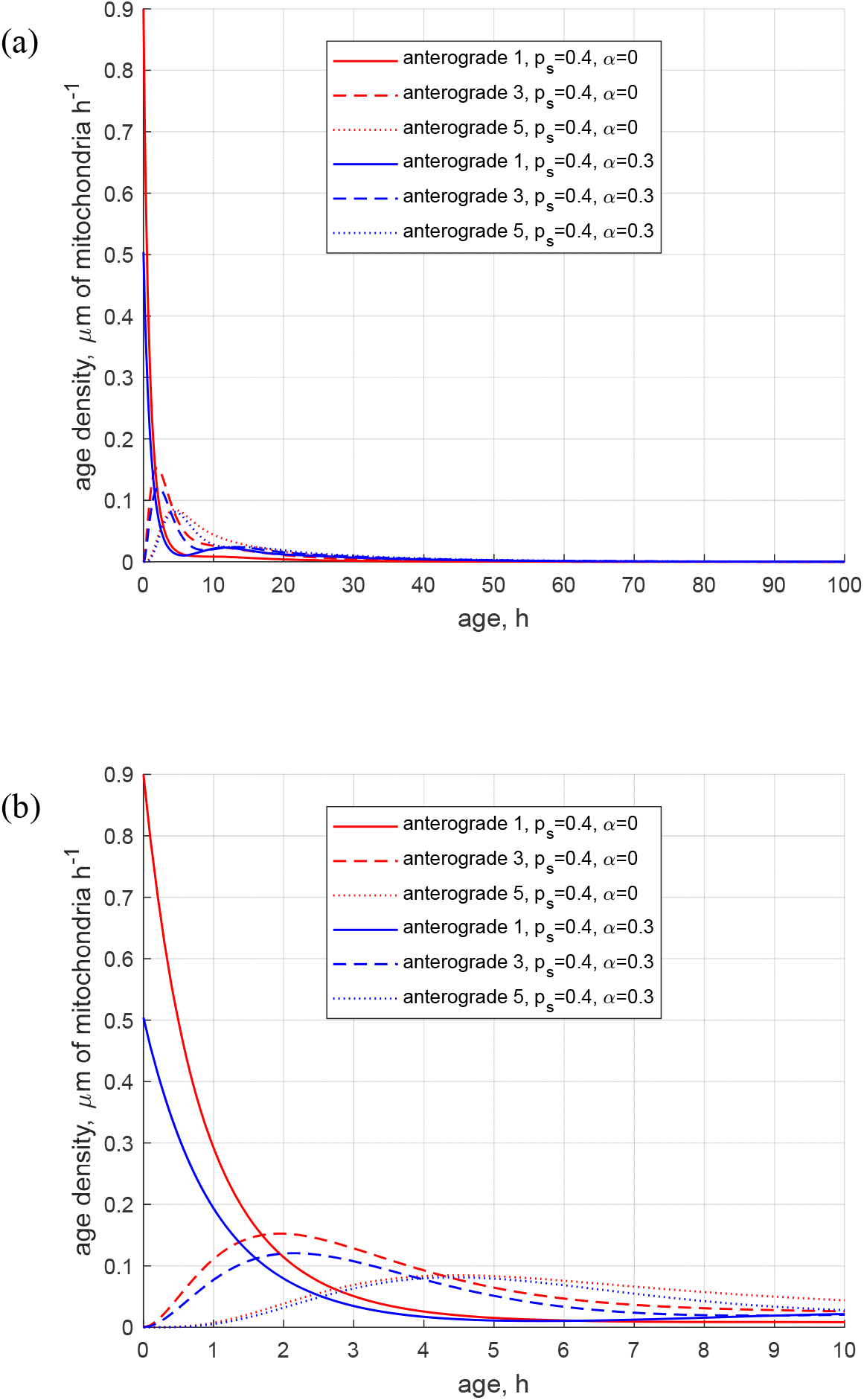
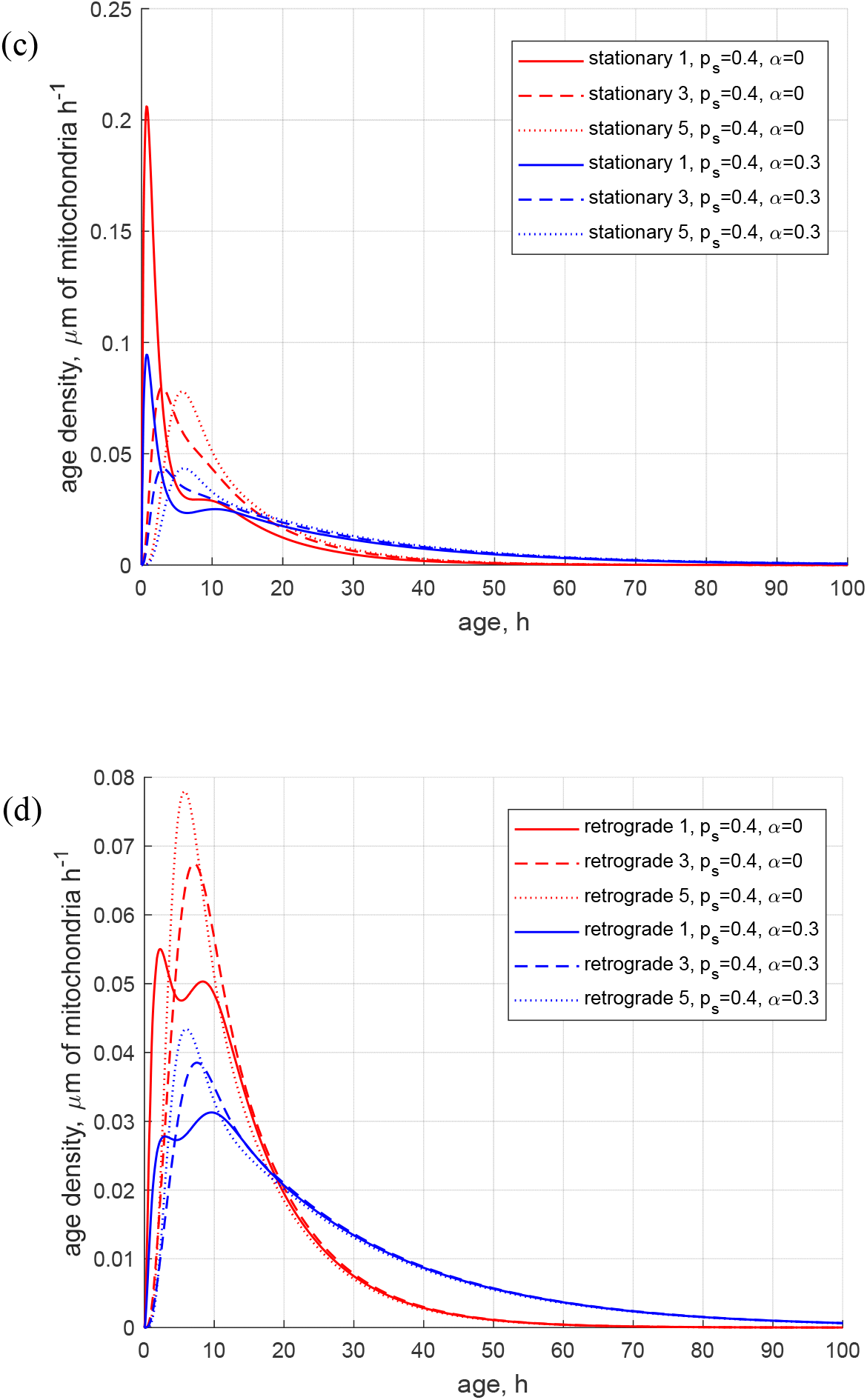
(a) Age density of anterogradely moving mitochondria in various demand sites. (b) Similar to Fig. S4a, this graph magnifies a specific age range of [0 10 hours] of the *x*-axis. (c) Age density of mitochondria in the stationary state in various demand sites. (d) Age density of retrogradely moving mitochondria in various demand sites. Two sets of parameter values: *p*_*s*_ = 0.4, *α* = 0 (base case) and *p*_*s*_ = 0.4, *α* = 0.3. Number of demand sites *N*=5.

**Fig. S5.**
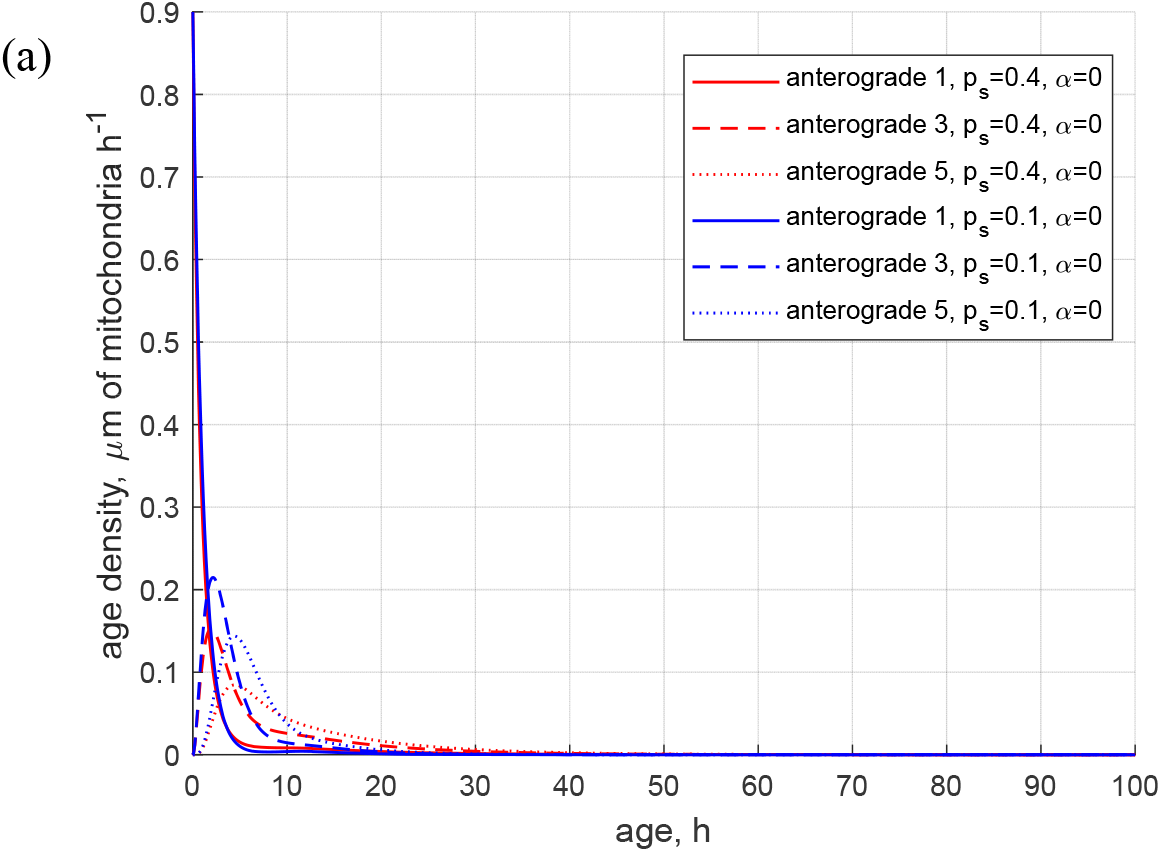
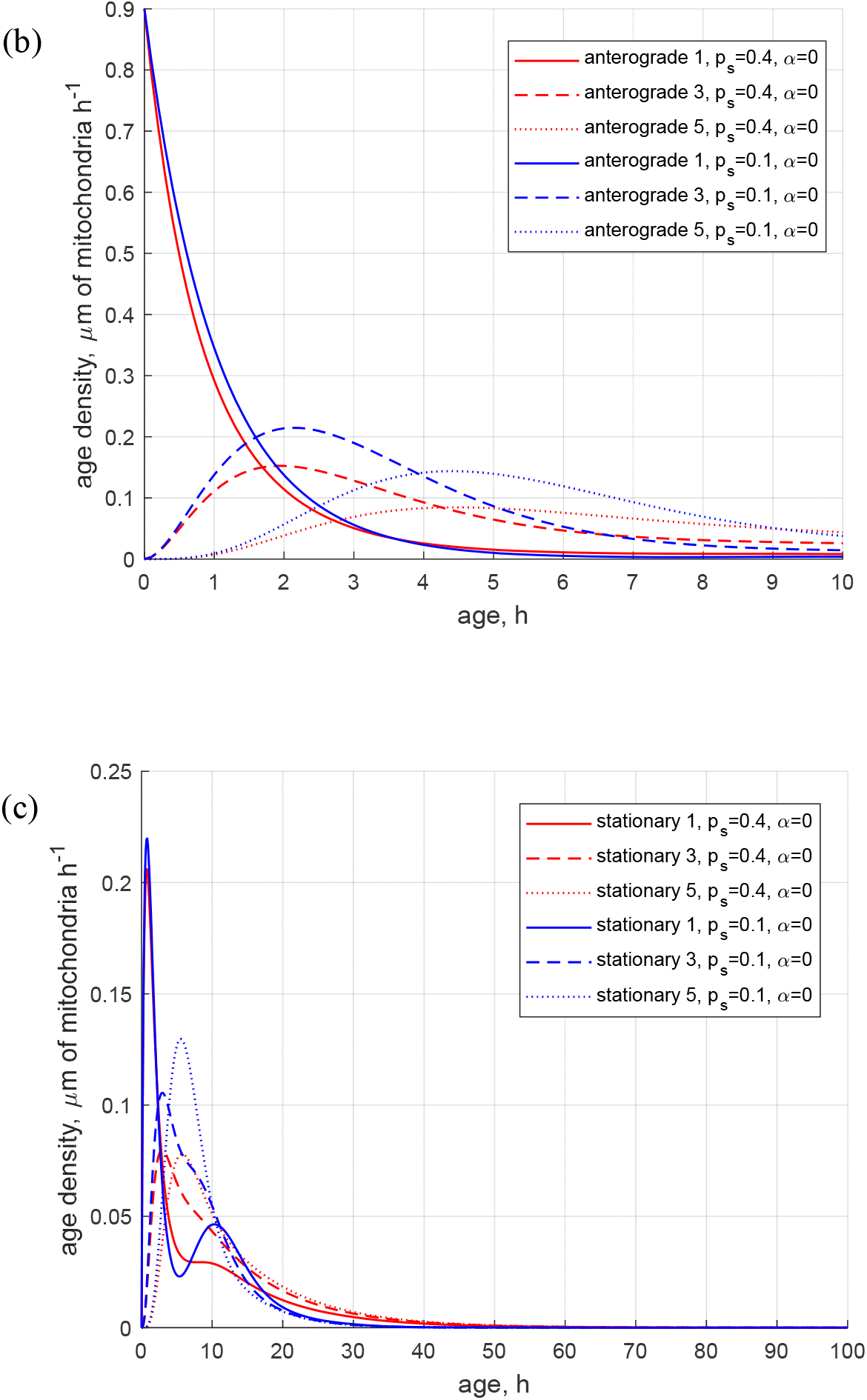
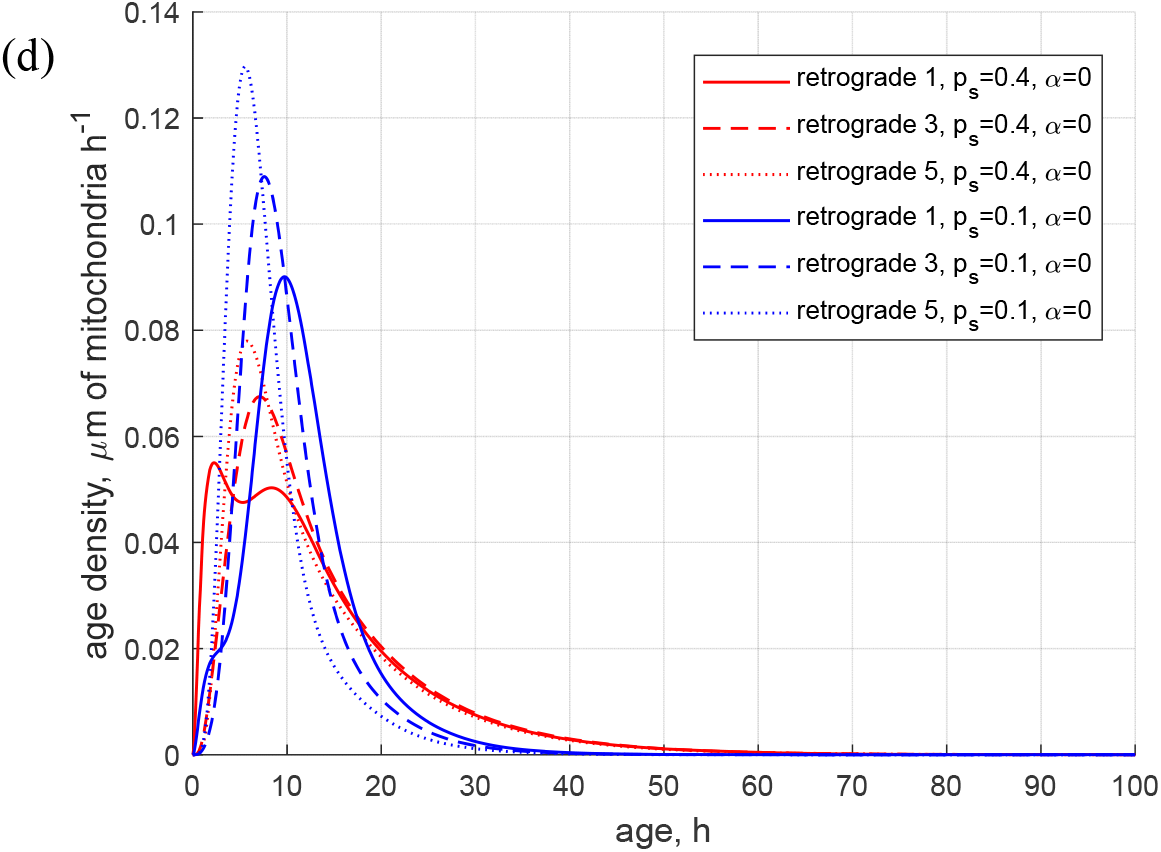
(a) Age density of anterogradely moving mitochondria in various demand sites. (b) Similar to Fig. S5a, this graph magnifies a specific age range of [0 10 hours] of the *x*-axis. (c) Age density of mitochondria in the stationary state in various demand sites. (d) Age density of retrogradely moving mitochondria in various demand sites. Two sets of parameter values: *p*_*s*_ = 0.4, *α* = 0 (base case) and *p*_*s*_ = 0.1, *α* = 0. Number of demand sites *N*=5.

**Fig. S6.**
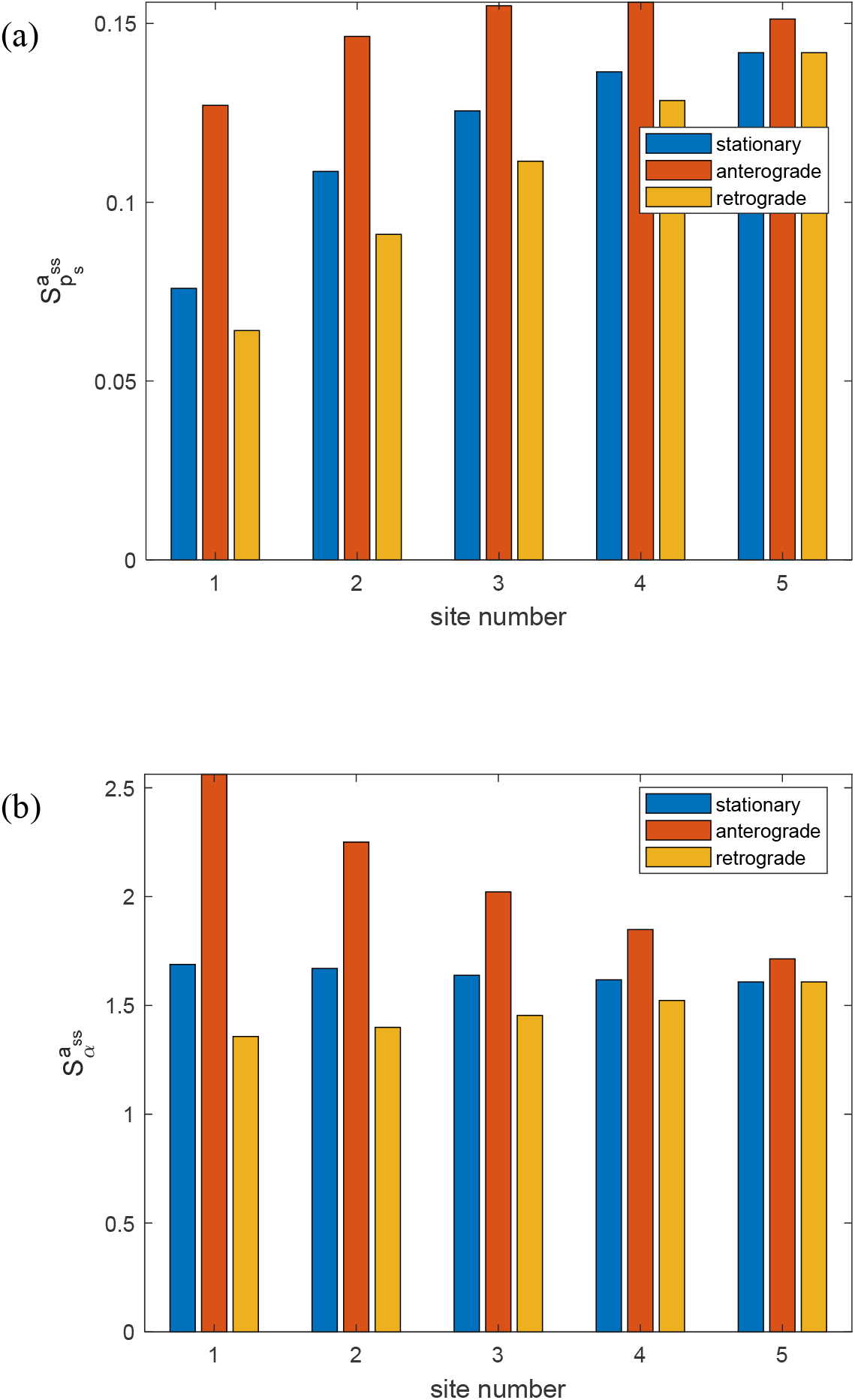
(a) Dimensionless sensitivity of the mean age of mitochondria to the probability that mobile mitochondria would transition to a stationary state at a demand site, *p*_*s*_, versus the site number. Computations were performed with Δ*p*_*s*_ = 10^−1^ *p*_*s*_. A close result was obtained for Δ*p*_*s*_ = 10^−2^ *p*_*s*_. (b) Dimensionless sensitivity of the mean age of mitochondria to the portion of mitochondria that return to the axon after exiting the axon, *α*, versus the site number. Computations were performed with Δ*α* = 10^−1^*α*. A close result was obtained for Δ*α* = 10^−2^*α*. Sensitivities are analyzed around *p*_*s*_ = 0.1 s^-1^, *α* = 0.3. Number of demand sites *N*=5.

## Notes

### Competing Interest Statement

The authors have declared no competing interest.

### Summary of Updates

We added a statement that the sensitivity of the mean age of stationary mitochondria to the stopping probability increases proportionally with the number of compartments in the axon. This suggests that in an extremely long axon with a large number of compartments, the age of mitochondria could be highly influenced by the stopping probability.

